# Automating parameter selection to avoid implausible biological pathway models

**DOI:** 10.1101/845834

**Authors:** Chris S. Magnano, Anthony Gitter

**Author notes:** Send correspondence to AG.

## Abstract

A common way to integrate and analyze large amounts of biological “omic” data is through pathway reconstruction: using condition-specific omic data to create a subnetwork of a generic background network that represents some process or cellular state. A challenge in pathway reconstruction is that adjusting pathway reconstruction algorithms’ parameters produces pathways with drastically different topological properties and biological interpretations. Due to the exploratory nature of pathway reconstruction, there is no ground truth for direct evaluation, so parameter tuning methods typically used in statistics and machine learning are inapplicable. We developed the pathway parameter advising algorithm to tune pathway reconstruction algorithms to minimize biologically implausible predictions. We leverage background knowledge in pathway databases to select pathways whose high-level structure resembles that of manually curated biological pathways. At the core of this method is a graphlet decomposition metric, which measures topological similarity to curated biological pathways. In order to evaluate pathway parameter advising, we compare its performance in avoiding implausible networks and reconstructing pathways from the NetPath database with other parameter selection methods across four pathway reconstruction algorithms. We also demonstrate how pathway parameter advising can guide construction of an influenza host factor network. Pathway parameter advising is method-agnostic; it is applicable to any pathway reconstruction algorithm with tunable parameters. Our pathway parameter advising software is available on GitHub at https://github.com/gitter-lab/pathway-parameter-advising and PyPI at https://pypi.org/project/pathwayParameterAdvising/.

## 1 Introduction

Network analysis can integrate and analyze large amounts of biological “omic” data from genomic, transcriptomic, proteomic, or metabolomic assays (Goh *et al*., 2012; Furlong, 2013). Placing omic data in a network context allows for the discovery of key members of a process that may be missed from a single data source and functional summarization for hypothesis generation and other downstream analyses.

Although biological pathway enrichment can be used to interpret omic data, pathways in curated databases are incomplete and also contain proteins or genes that are not involved in a particular biological context (Köksal *et al*., 2018). Thus, it is often preferable to infer a customized subnetwork specific to an experimental dataset starting from all known protein interactions, referred to as the interactome. We refer to this problem as pathway reconstruction: using condition-specific input omic data to select nodes and edges that represent some process or cellular state from a generic background network. Pathway reconstruction differs from module detection (Choobdar *et al*., 2019), which divides a network into functional units or clusters. It is also distinct from network propagation (Cowen *et al*., 2017), which identifies relevant regions of a larger network but typically does not select specific edges within that region.

Existing pathway reconstruction methods are based on diverse strategies such as combinatorial optimization problems (Tuncbag *et al*., 2016; Scott *et al*., 2006; Yosef *et al*., 2011), shortest paths (Ritz *et al*., 2016), enrichment analysis (Cerami *et al*., 2010), network flow (Basha *et al*., 2013; Goldberg and Tarjan, 1990; Komurov *et al*., 2010), and other graph theory algorithms. These methods also take in a variety of inputs. Some, such as the Prize-Collecting Steiner Forest (PCSF) algorithm (Tuncbag *et al*., 2016; Kedaigle and Fraenkel, 2018), accept scores for the biological entities of interest. Other methods, such as PathLinker (Ritz *et al*., 2016), require the inputs to be split into starting-points (sources) and end-points (targets) for the pathway. Despite the different optimization strategies and inputs, pathway reconstruction algorithms almost always require the user to set parameters. Adjusting the parameters can produce pathways with drastically different topological properties and biological interpretations. For instance, both pathways in Figure 1 were created with the same PCSF algorithm and the same influenza host factor screen data (Section 4.6.3); they only differ in the PCSF parameters used. The pathway on the right is reasonably sized and can be interpreted and summarized for downstream analysis. The pathway on the left, however, includes over 7,000 nodes and would impractical to interpret or analyze. We refer to these types of pathways, pathways that are either impractical to analyze or contain topological features that are biologically unrealistic, as being *implausible*.

**Fig. 1.**
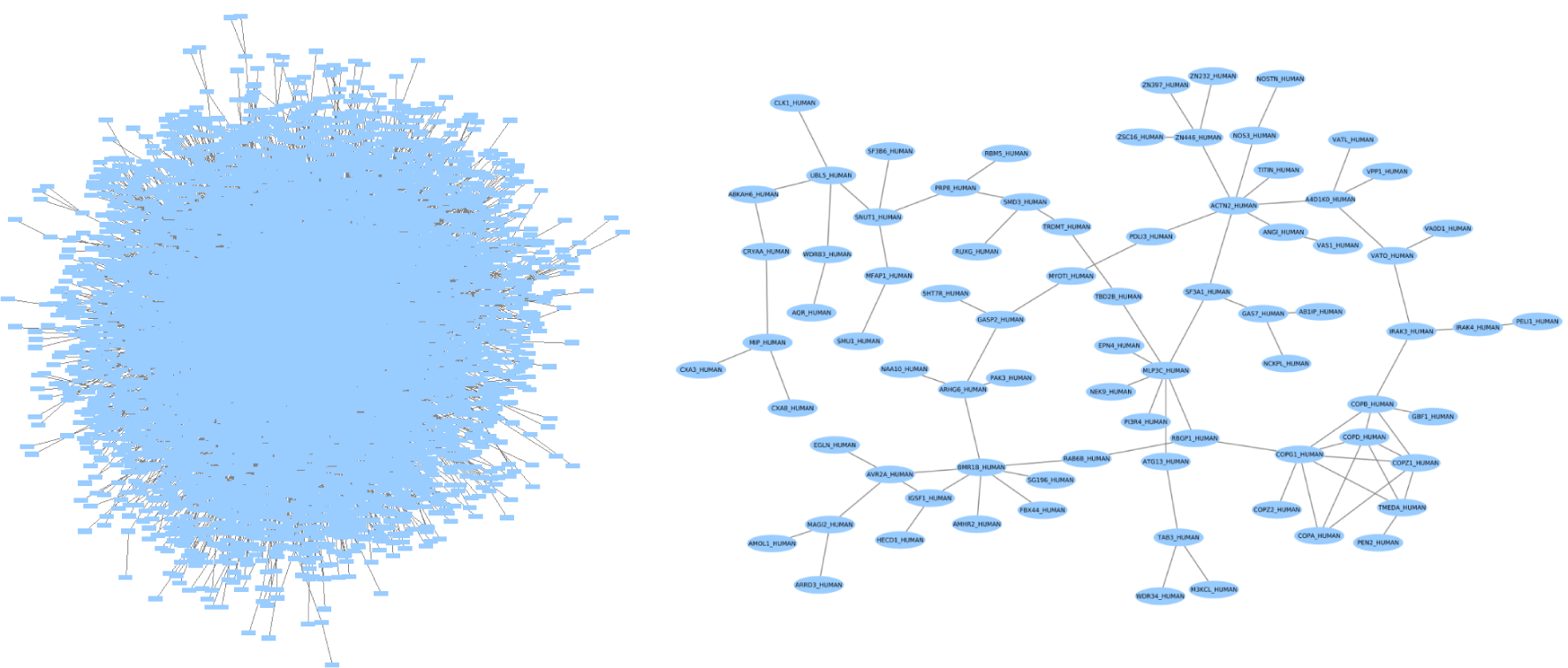
Influenza host factor pathways created using PCSF from RNA interference (RNAi) screens (Section 2.4), here showing the largest connected components from ensembling the bottom 100 ranked pathways (left) and the top 100 ranked pathways (right). The only difference in creating the networks was the range of PCSF parameter values.

An open challenge is how to configure these critical pathway reconstruction parameters in a manner that is objective and applicable across diverse types of algorithms. Existing approaches tend to be informal and ad hoc, and most best practices in parameter tuning are not applicable to pathway reconstruction. The simplest way to choose a parameters would be to use default values. However, a single parameter setting cannot work for all datasets. The number of input proteins, genes, or metabolites varies based on the experiment, and the effects of input size can be unpredictable for different pathway reconstruction algorithms. For instance, if PathLinker is run with fixed parameters, increasing the number of source and target nodes can actually result in a smaller reconstructed pathway, which is not necessarily what a user would intend. In addition, some methods are commonly run repeatedly and combined into an ensemble network (MacGilvray *et al*., 2018; Budak *et al*., 2015; Khurana *et al*., 2017), which requires multiple parameter settings.

Parameter tuning methods from supervised learning are also a poor match. There is no ground truth for supervised parameter tuning, and unsupervised cross-validation (CV) is ineffective (Section 2). In addition, the objective functions of pathway reconstruction methods only approximate biologically meaningful graph topologies and typically have no probabilistic likelihood. Thus, their values cannot be compared between different parameter settings, and statistical model selection criteria such as the Akaike information criterion (Akaike, 1998) and the Bayesian information criterion (Schwarz, 1978) are not applicable. For instance, the PCSF objective function value can be arbitrarily increased by changing the parameter values.

Given the lack of objective, quantitative methods for tuning, parameter settings are often chosen by manual inspection or informal heuristics. ResponseNet recommends choosing parameters that recover at least 30% of inputs while minimizing low confidence edges (Yeger-Lotem *et al*., 2009). PCSF recommends choosing pathways robust to small random input variation or matching the average degree of input nodes and non-input nodes (Kedaigle and Fraenkel, 2018). We show in Section 2 that these heuristics can perform poorly in practice.

Biologists often have intuition about which pathways are unrealistic or impractical for downstream analysis, such as the 7,000 node pathway in Figure 1 or subnetworks with unusual degree distributions. Pathway reconstruction is typically an exploratory analysis used to summarize the input data and generate hypotheses leading to further experiments. In this context, it is important to avoid implausible and uninterpretable pathway topologies. Therefore, parameter tuning should not focus on traditional notions of accuracy but instead formalize how *useful* reconstructed pathways are to biologists. Our major contribution is providing a formal approach based on graph topology that quantifies this biological intuition and can be used to optimize pathway reconstruction in an objective manner.

One framework for finding optimal parameters in an uncertain setting is parameter advising (Kececioglu and DeBlasio, 2013; DeBlasio and Kececioglu, 2015, 2017), which was originally developed for multiple sequence alignment. Parameter advising can be used to adapt the parameter tuning framework in settings where no ground truth tuning set exists. However, parameter advising requires a means to estimate the accuracy of a model with a given set of parameters. Because pathway reconstruction is an exploratory analysis, there is no formal notion of accuracy. We overcome this limitation by leveraging background knowledge to create a score that prefers pathways in topological agreement with reference pathways. Our parameter tuning method, pathway parameter advising, uses the parameter advising framework in combination with a distance metric based on graphlet decomposition to measure similarity between reconstructed pathways and pathways from curated databases. Pathway databases may be imperfect and incomplete, but they reflect models that the expert curators consider to be biologically plausible. Only measuring the topology of a reconstructed pathway means that pathway parameter advising is also method agnostic. It can tune the parameters of any pathway reconstruction method.

Figure 2 gives an overview of the experiments performed to evaluate pathway parameter advising. We explore the effectiveness of pathway parameter advising in avoiding parameters that lead to implausible networks by reconstructing 15 pathways from the NetPath database. Pathway parameter advising outperforms other parameter selection methods and default parameter values. We also examine how well these reconstructed pathways overlap with the original NetPath pathways. Finally, we show how pathway parameter advising can guide pathway reconstruction with an influenza host factor network.

**Fig. 2.**
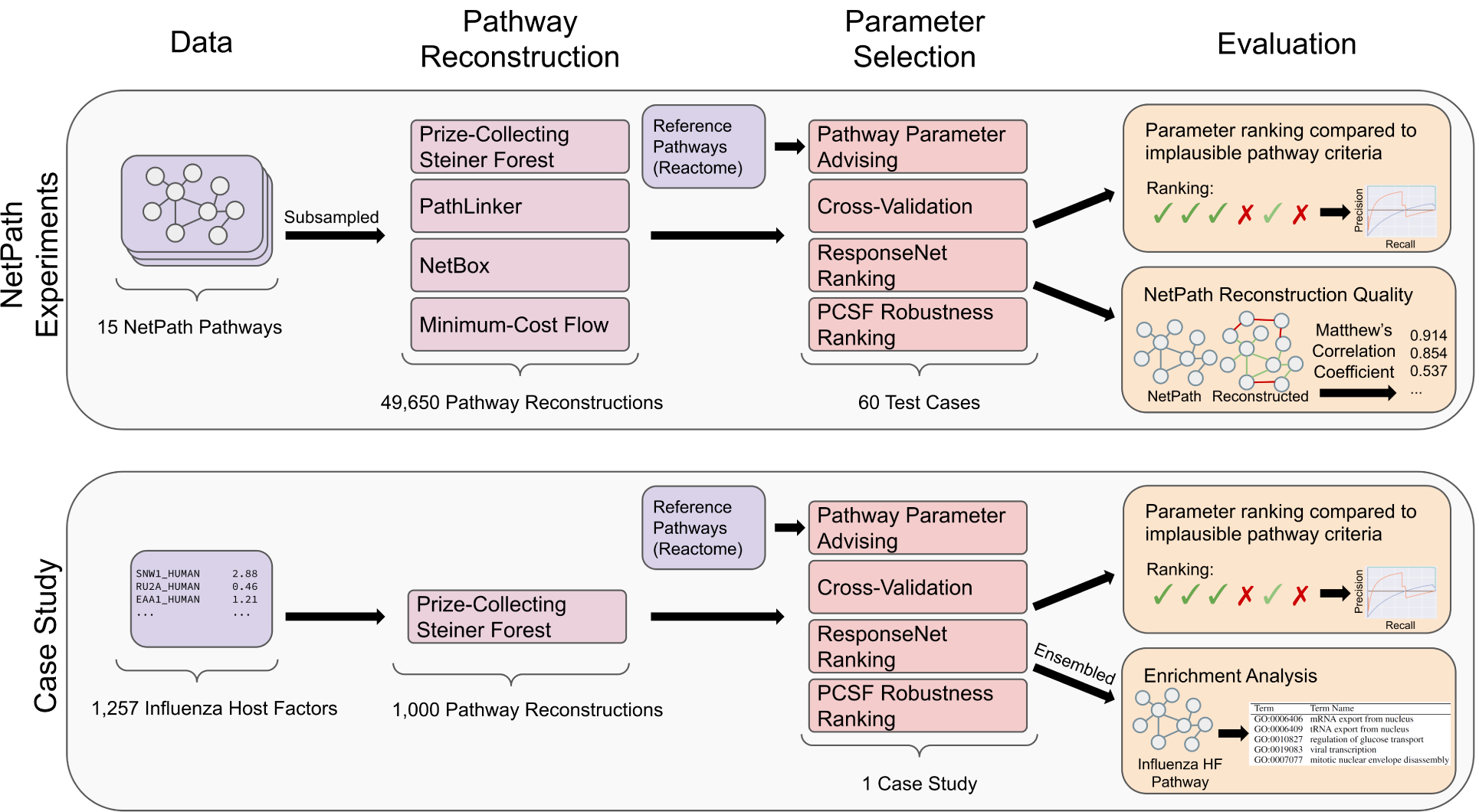
Overview of experiments performed to evaluate pathway parameter advising.

## 2 Results

Pathway parameter advising ranks parameter settings for pathway reconstruction methods. Because pathway reconstruction is an exploratory analysis and our goal is to maximize downstream biological utility, which cannot be directly quantified, we resort to multiple indirect approaches to evaluate pathway parameter advising (Figure 2). We first use the literature to define topological graph properties that make a candidate pathway biologically implausible. Pathway parameter advising can optimize pathway reconstruction algorithms to avoid parameters that lead to implausible pathways. Next, we demonstrate that we can improve the reconstruction of NetPath pathways by comparing predicted pathway edges with the NetPath ground truth. We then show how pathway parameter advising can be applied in practice and reconstruct an influenza host factor pathway from RNAi screens. Finally, we ensure that the graphlet distance ranking metric at the core of pathway parameter advising has desirable properties. It is not overly sensitive to disconnected graphlets and shows that reference pathways are similar to one another.

### 2.1 Pathway Parameter Advising Overview

Pathway parameter advising is a framework to score the pathways produced by a pathway reconstruction algorithm run with multiple parameter settings. The scored pathways are ranked so that the top-scoring pathway(s) can be used for downstream analyses and poor-scoring pathways can be ignored. We define a pathway quality score for the ranking that compares the graph structure of a reconstructed pathway with the graph structures found in reference pathways from a curated database. Specifically, the score computes the topological distances between the reconstructed pathway and reference pathways. Reconstructed pathways with small distances are preferred.

To compute the pathway quality score, the reconstructed pathway is first decomposed into small subgraphs called graphlets (Pržulj *et al*., 2004). We calculate the graphlet frequencies in the reconstructed pathway and the reference pathways. This graphlet frequency decomposition summarizes the topology of a pathway in a vector of 17 values, one per graphlet type (Figure 3). The distances between the reconstructed pathway and the reference pathways are calculated by computing the difference between these graphlet decomposition vectors. The final metric used to rank parameter settings is defined as the mean distance between the reconstructed pathway and the 20% closest reference pathways. More details on the pathway parameter advising algorithm can be found in Section 4.1.

**Fig. 3.**
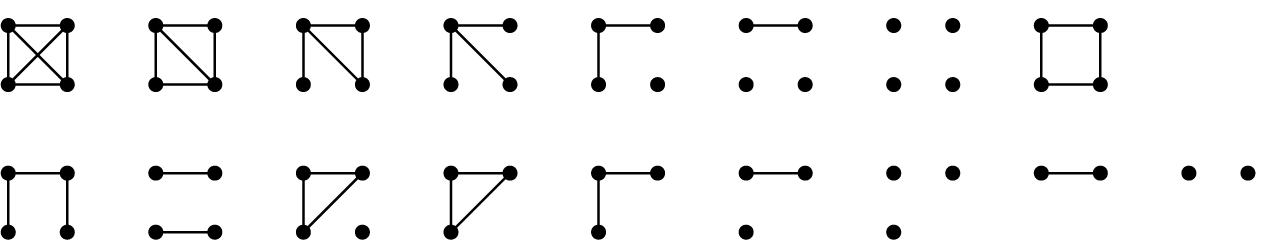
Pathways are decomposed into these 17 graphlets for graphlet frequency distance calculations.

### 2.2 Implausible Pathway Detection

In order to evaluate pathway parameter advising, we considered its ability to avoid implausible networks. While it is difficult to define a single best pathway in the context of an exploratory analysis, some pathways are clearly biologically unrealistic, infeasible to analyze, or not useful for downstream analysis. Thus, pathway parameter advising should consistently rank parameter settings that lead to plausible networks above those that lead to implausible networks.

We applied 4 pathway reconstruction methods to sampled NetPath pathways (Section 4.6) to reconstruct the 15 NetPath pathways. We selected the NetPath pathways Wnt, TNF alpha, and TGF beta as validation pathways to evaluate different graphlet distances and develop pathway parameter advising. These 3 validation pathways are excluded from all aggregate results. The other 12 NetPath pathways were designated as test pathways. More details on the 4 pathway reconstruction methods, NetBox, PathLinker, PCSF, and min-cost flow, can be found in Section 4.2. Overall, we considered 60 parameter tuning tasks across the 15 NetPath pathways and 4 pathway reconstruction methods.

In these experiments, we used the implausibility criteria defined in Section 4.4 to label reconstructed pathways as plausible or implausible without reference to Reactome or other reference pathways. Reconstructed pathways that meet all of the plausibility criteria are treated as the positive set, while reconstructed pathways defined as implausible are the negative set. These reconstructed pathway labels of plausible and implausible are considered alongside the ranking produced by a parameter selection method. We then used precision-recall (PR) curves (Figures S1-S5) to evaluate how well pathway parameter advising and alternative parameter selection strategies distinguish plausible from implausible pathways. Parameter selection methods that rank plausible pathways above implausible pathways will have a higher area under the PR curve (AUPR).

We compared pathway parameter advising to three other parameter selection methods: CV, ResponseNet ranking, and PCSF robustness ranking (Section 4.3). Figure 4 shows the distribution of AUPRs across the 4 pathway reconstruction methods on the NetPath test pathways. Different pathway reconstruction methods had varying proportions of networks identified as plausible, with min-cost flow having the lowest mean proportion at 11% and NetBox with the highest at 89%. Among the 12 test pathways, pathway parameter advising had the highest median AUPR for each pathway reconstruction method.

**Fig. 4.**
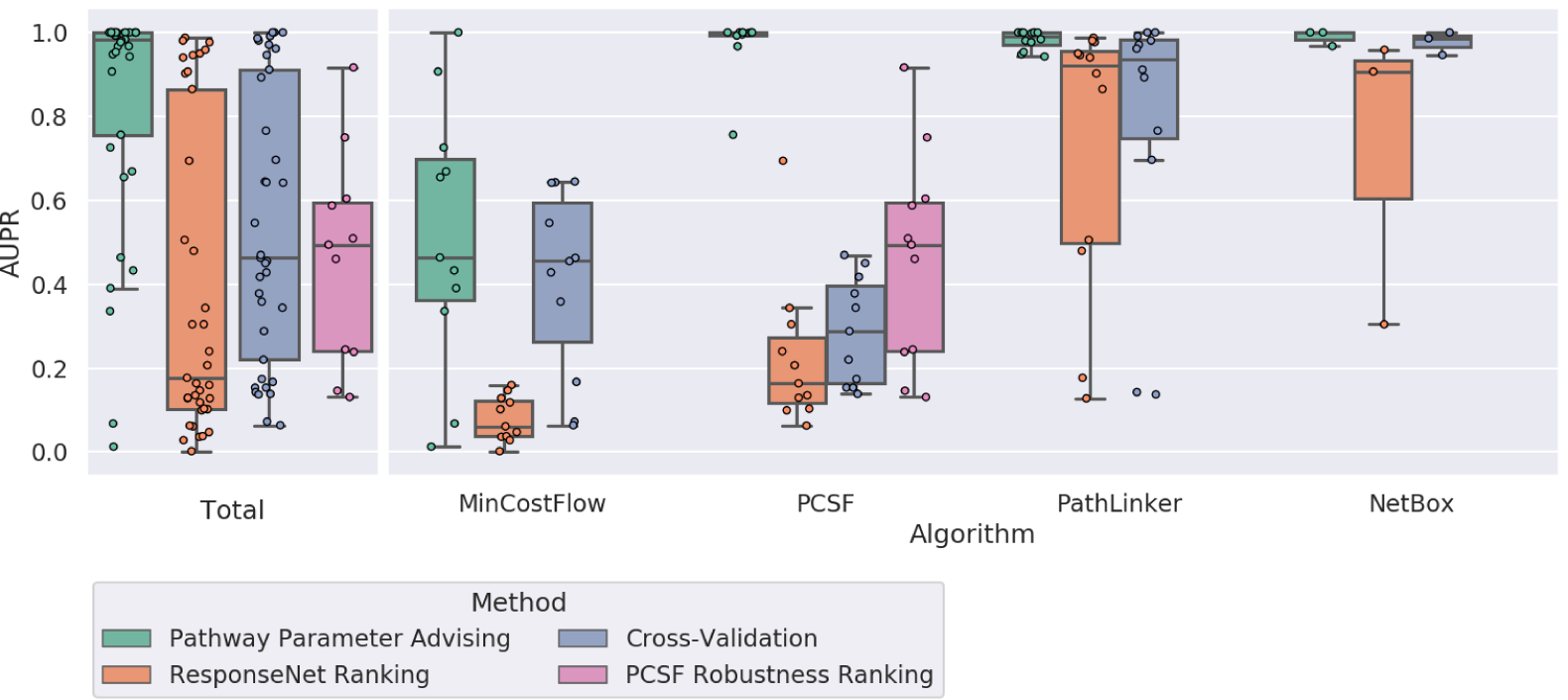
Performance of parameter selection methods on avoiding implausible networks. Boxplots represent the distributions of the AUPRs aggregated for 4 pathway reconstruction methods and 12 NetPath test pathways. Degenerate cases where all or no pathways met the plausibility criteria are excluded. Full results, including the 3 validation pathways and degenerate cases, can be found in Figures S2-S5.

Of the 36 cases where AUPRs could be compared (both plausible and implausible pathways were present), pathway parameter advising had the highest AUPR in 30. CV had the highest AUPR in the other 6. The impact of the choice of parameter ranking strategy is most stark for PCSF, where pathway parameter advising has perfect AUPR in almost all pathways and the other approaches struggle. Not only did pathway parameter advising have the highest median AUPR overall, but its performance was the most consistent; it had the lowest variance in AUPR across all tasks.

In order to make sure that performance was not overly influenced by our specific choice of criteria for plausible and implausible pathways, we varied the threshold for each of the 4 plausibility topological properties. We considered all combinations of thresholds for all 4 proprieties in a grid search, resulting in a total of 10,000 plausibility configurations. Figure 5 shows the AUPR of each parameter ranking method as the plausibility threshold for each topological feature is varied. Figure S1 shows aggregate AUPR values. Pathway parameter advising outperforms all other parameter ranking methods at each threshold value.

**Fig. 5.**
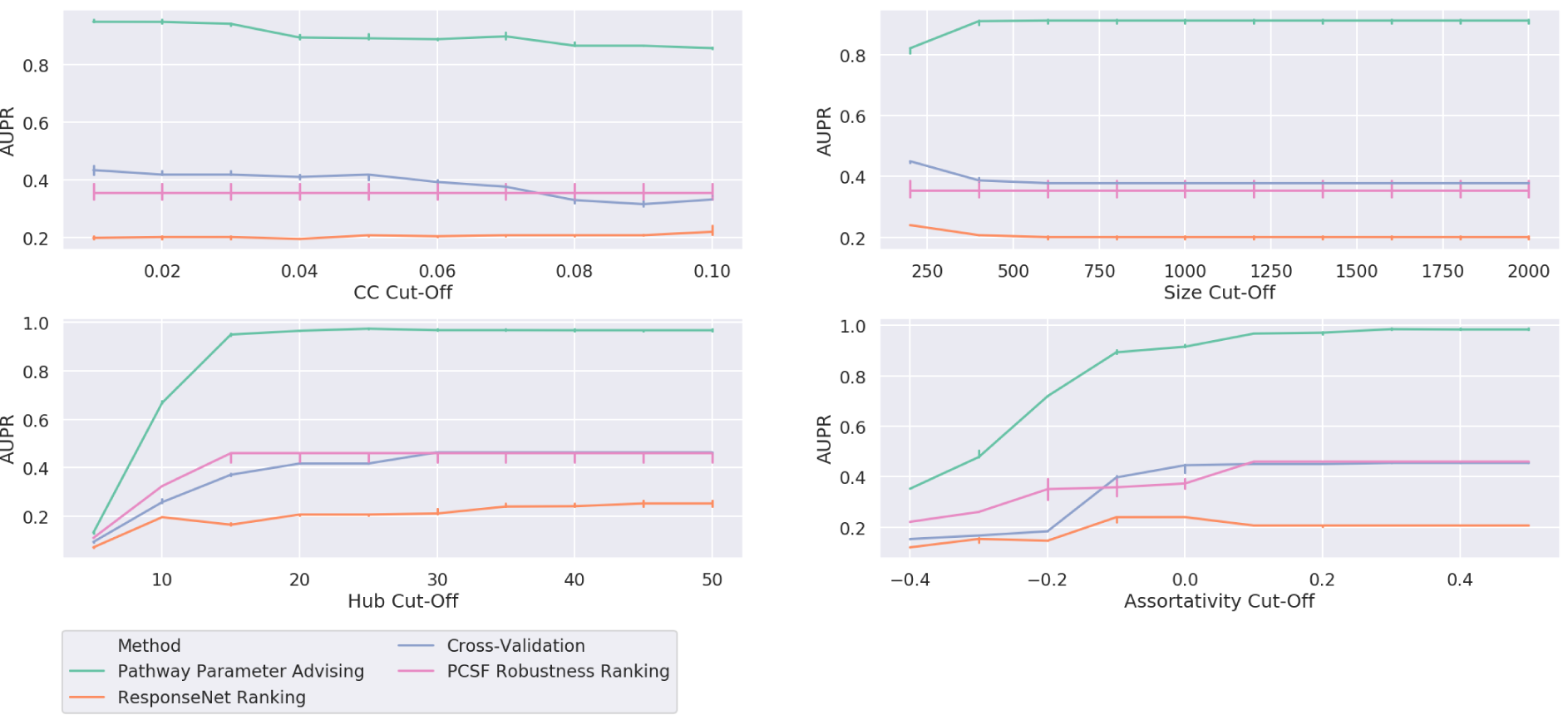
Performance of parameter selection methods on avoiding implausible networks as the threshold for plausibility is varied across different topological features – clustering coefficient, pathway size, hub node dependence, and assortativity – as described in Section 4.4. Lines show median AUPR over the varied thresholds for the other 3 topological features for all 12 NetPath test pathways and 4 pathway reconstruction methods. Error bars show the 95% confidence interval.

### 2.3 Quality of NetPath Pathway Reconstruction

Having achieved our primarily goal of accurately prioritizing parameters that generate plausible pathways, we also evaluated the quality of the pathway reconstructions themselves. We compared pathway parameter advising to the alternative ranking methods and the default parameters. In these experiments, we define all pathway edges in the NetPath pathway as a positive set and all other edges as a negative set. We then compared the ability of pathway parameter advising and other parameter ranking methods to promote reconstructed pathways that closely resemble their NetPath equivalent.

Figure 6 (left) shows the adjusted Matthew’ s Correlation Coefficients (MCCs) of all 48 pathway reconstruction tasks. MCC quantifies the overlap between the reconstructed pathway edges and NetPath pathway edges (Section 4.5). While pathway parameter advising has the highest median adjusted MCC, the parameter selection method has less impact on MCC than it did on pathway plausibility (Figure 4). When stratified by pathway reconstruction algorithm, pathway parameter advising has the highest median adjusted MCC for PCSF and PathLinker. However, the default parameters perform almost as well for PathLinker. CV has the highest adjusted MCC in min-cost flow and NetBox. Of the 48 pathway reconstruction tasks, pathway parameter advising had the highest median adjusted MCC 21 times, while CV had 13, default parameters had 9, and the ResponseNet ranking had 8, including 3 cases where 2 methods tied.

**Fig. 6.**
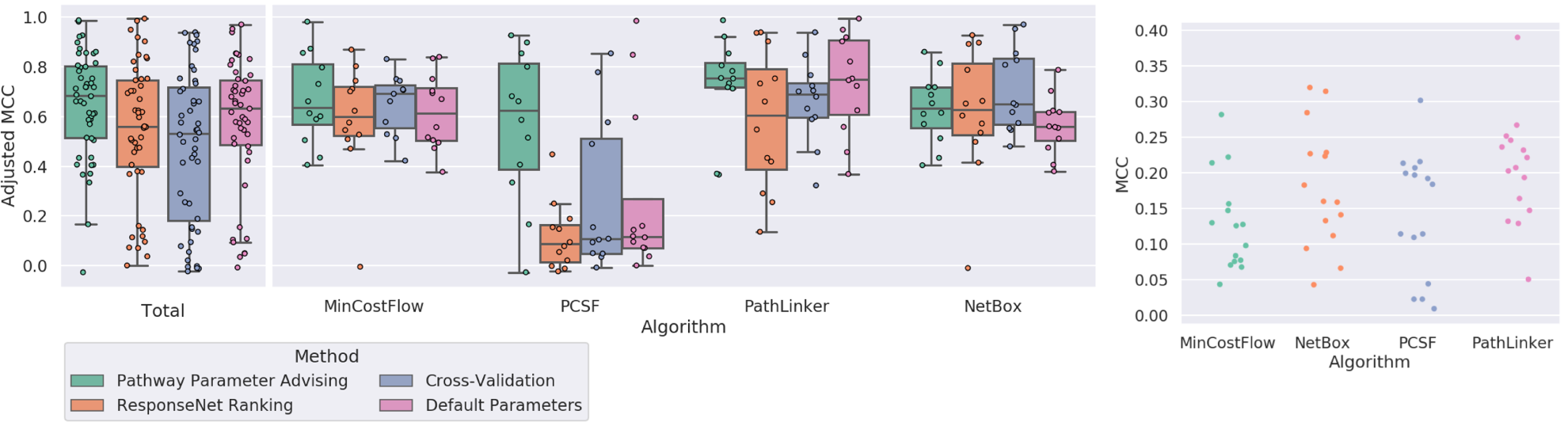
Left: Adjusted MCC of parameter selection methods on reconstructing 12 NetPath test pathways across 4 pathway reconstruction methods. MCCs were adjusted by normalizing them to the highest possible MCC within a given pathway reconstruction method and pathway. Figure S6 shows MCC values by pathway, including the 3 validation pathways excluded here. Right: The highest possible MCC of pathway reconstruction in 60 parameter sweeps across 4 pathway reconstruction methods and 15 NetPath pathways (validation and test). The MCC values are generally low, reflecting low overlap between the predicted and NetPath pathway edges.

Figure 6 (right) shows the distribution of best possible unadjusted MCCs for all pathway reconstruction tasks, including the 3 NetPath validation pathways. The best possible MCC was greater than 0.3 in only 4 of the 60 cases, and it was never greater than 0.4. Given these low unadjusted MCC values and the overall objective of our study, the implausible pathway detection experiment is a better indicator of parameter selection performance.

### 2.4 Influenza Host Factor Pathway Reconstruction

To illustrate how pathway parameter advising can guide the biological interpretation of omic data, we reconstructed an influenza host factor pathway. Our aim was to create a pathway that represents aspects of influenza’ s infectious activities and could lead to the discovery of new host factors or host factor regulators. We created an influenza host factor network using the 1,257 host factors from a meta-analysis of 8 RNAi screens (Tripathi *et al*., 2015). These host factors were given as input to PCSF using the same range of parameter settings as in the other experiments, creating a candidate host factor pathway for each parameter setting. We used the magnitude of the consolidated Z scores given in the meta-analysis as node scores (see Section 4.6.3).

After creating the candidate host factor pathways, we ranked the 1,000 parameter settings using pathway parameter advising. Figure 7 (left) shows the PR curve of different parameter ranking methods’ ability to avoid implausible networks. Pathway parameter advising ranked the pathways almost perfectly, with an AUPR of 0.96, while other parameter selection methods had more difficulty separating implausible from plausible networks. CV and the ReponseNet rankings performed worse than a random ranking. Pathway parameter advising performs well not only on simulated data from NetPath but also on data aggregated from real high-throughput experiments.

**Fig. 7.**
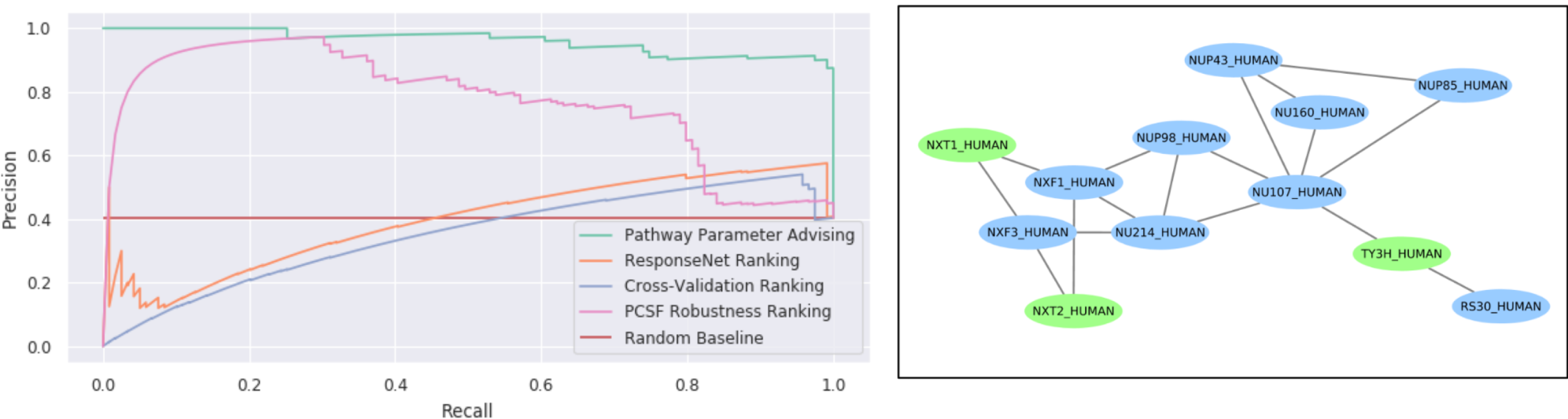
Left: Precision-recall curve for implausible networks in PCSF influenza host factor network construction. Right: A component of the influenza host factor ensemble pathway created from the top 50 PCSF parameter settings ranked by pathway parameter advising. This component represents 12 of the 86 total nodes in the pathway (Figure S7). Host factor nodes provided as input are shown in blue, while green nodes are “Steiner” nodes that PCSF predicts to connect the host factors.

We also created three ensemble networks from the resultant pathways from the top, middle, and bottom 50 parameter settings ranked by pathway parameter advising. As discussed in Section 1, ensembling the reconstructed pathways is a common way to use PCSF. We expect the top 50 ensemble pathway to be best for downstream analysis and interpreting the input host factor data. Of the 3 ensemble pathways, the ensemble made from the 50 highest ranked parameters contains 86 nodes. The middle ranked and low ranked ensemble pathways have 7,337 and 15 nodes, respectively (Figure S7). The middle ranked pathway is too large to interpret and does not provide meaningful new insights into the relationships among host factors. The low ranked pathway is too small to illuminate new biological hypotheses. The top ranked pathway, however, is large enough for meaningful enrichment and downstream analyses while remaining small enough to be feasible.

We then performed a gene set enrichment analysis on the top ensemble pathway using DAVID (Huang *et al*., 2008) (Section 4.7). We tested both Gene Ontology (GO) biological process terms (Table S1) and Kyoto Encyclopedia of Genes and Genomes (KEGG) pathway enrichment (Table S2). The top 2 KEGG pathways enriched were RNA transport and influenza A. The influenza A pathway being enriched is a confirmation that our ensemble pathway represents influenza processes well.

The RNA transport pathway enrichment captures one unique aspect of influenza A. It replicates within the nucleus, so it has complex processes for transporting viral RNA in and out of the nucleus (Dou *et al*., 2018). Similar concepts are observed in the top 2 enriched GO terms, mRNA and tRNA export from nucleus (Table S1). We also see other general viral GO terms, which confirm the top ranked pathway’ s representation of influenza, such as viral transcription and intracellular transport of virus. Viral transcription, mRNA export from nucleus, and tRNA export from nucleus all contain the same 6 ensemble pathway members, while mRNA export contained an additional 3 unique pathway members and viral transcription contained an additional 2 unique pathway members.

Figure 7 (right) shows one connected component representing 12 of the 86 nodes from the top ranked ensemble pathway. One node in particular, NXT2, was not among the original host factors but was identified as a possible host factor in a later genome-wide CRISPR/Cas9 screen (Han *et al*., 2018). This demonstrates how pathways chosen through pathway parameter advising could guide new discoveries.

The influenza study also illustrates the danger of running pathway reconstruction methods with default parameters alone. The PCSF network constructed using default parameters had 6,676 nodes, which is too large to interpret. We examined the role of NXT2 in the pathway reconstructed using default parameters. Figure 8 shows the subnetworks from this pathway for all nodes reachable within 3 and 2 edges of NXT2. As opposed to a functionally cohesive subnetwork, NXT2 only connects to a high-degree node. This node then connects to APP and SUMO2, which are the 2 highest degree nodes in the interactome. As a result, 727 nodes are within 3 edges of NXT2, which is more likely an artifact of the PCSF algorithm run with improper parameters than a meaningful biological pathway structure. This hub-based structure gives much less insight into the role of NXT2.

**Fig. 8.**
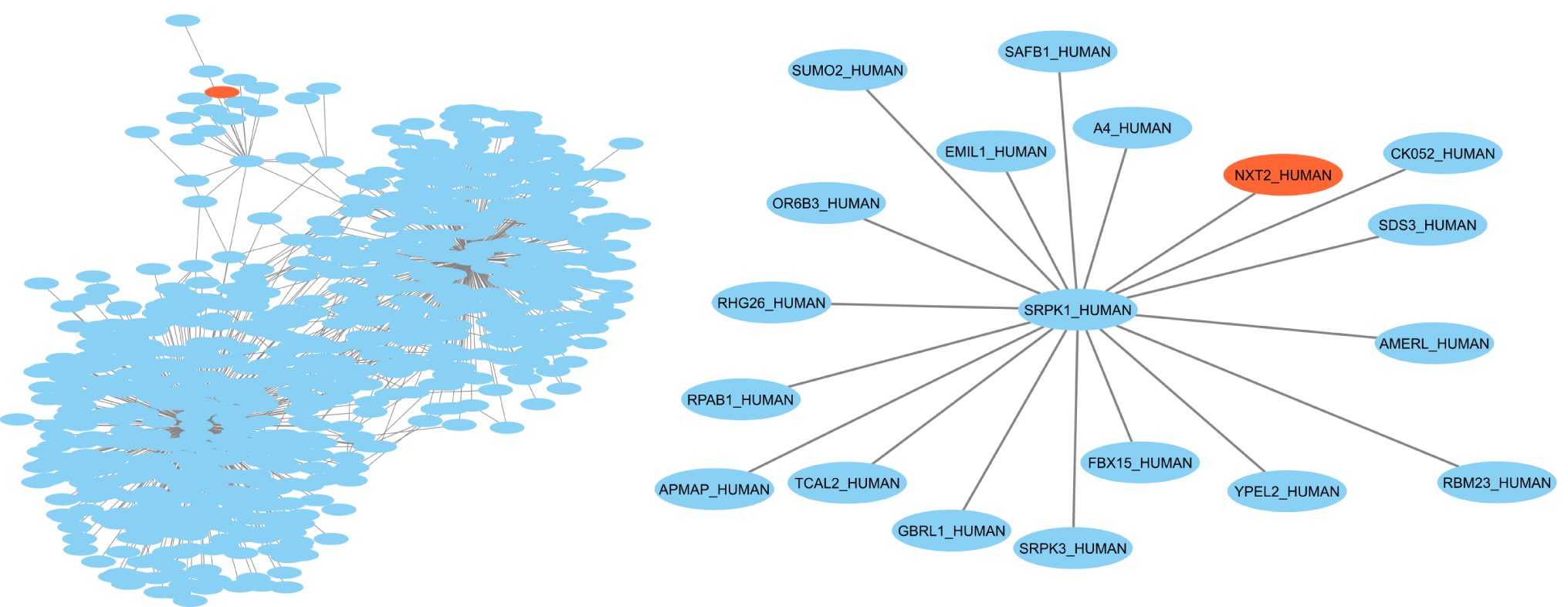
All nodes within distance 3 (left) and distance 2 (right) of NXT2, which is highlighted in orange, in the PCSF influenza host factor pathway reconstructed from default parameters. The default parameters resulted in a large hub-node focused pathway with little useful biological insight.

### 2.5 Evaluating the Ranking Metric

To further assess the validity of our pathway ranking metric, we explored several properties of the graphlet distance score. We first examined the overall distribution of graphlet distances (*E*(*G*) in Section 4.1.3) across all pathway reconstruction methods and Reactome pathways with added noise. *E*(*G*) is the score used by pathway parameter advising to rank pathways. We added noise to Reactome pathways by randomly removing a percentage of all edges in the pathway and adding back that number of edges that did not appear in the original pathway. We also calculated the graphlet distance between each Reactome pathway and all of the other Reactome pathways. Figure 9 shows these distributions of graphlet distances. Reactome pathways had a lower mean distance to other Reactome pathways than any set of reconstructed pathways, confirming that our metric ranks reference pathways over reconstructed pathways.

**Fig. 9.**
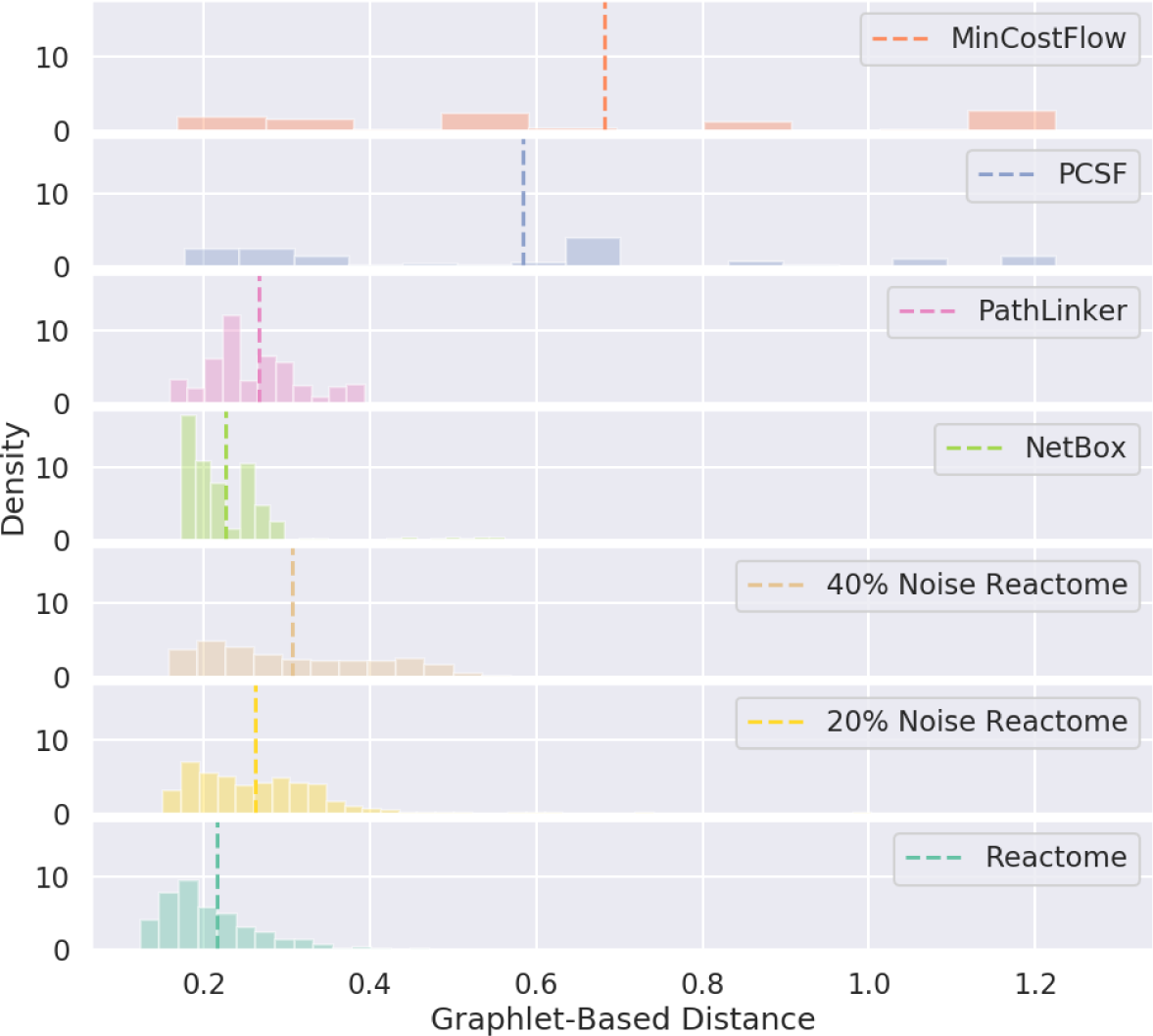
Distribution of aggregated graphlet-based distances (*E*(*G*)) for reconstructed pathways, Reactome pathways, and Reactome pathways with added noise. These aggregate distances were the metric used by pathway parameter advising to rank pathways, with lower distances being ranked higher. They were calculated by comparing the candidate pathway (reconstructed pathway, Reactome pathway, or noisy Reactome pathway) with all of the reference Reactome pathways. However, Reactome pathways were excluded from their own distance calculation. Vertical dashed lines show the mean graphlet distance.

We also explored how adding noise to the reference pathways affects the performance of pathway parameter advising. We assessed implausible pathways using Reactome pathways with increasing amounts of noise as the reference pathways. The level of noise had almost no effect on pathway parameter advising’ s ability to recognize and deprioritize implausible pathways (Figure S8). This may be due to how the noise randomly edited the Reactome pathways. If the random changes to the Reactome pathways are unbiased, then the changes in graphlet frequencies may cancel each other out in aggregate.

In order to examine the possibility that unconnected graphlets dominate the ranking metric, we first examined the breakdown of total graphlet distance by each of the individual graphlets in Figure 3. Figure S9 shows the contribution of each graphlet to the ranking metric. While a single graphlet, the 4 unconnected nodes graphlet, does have the largest contribution to graphlet distance, its median contribution to the ranking metric’ s value is only 30%. Because biological pathways tend to be sparse, this graphlet count scales with pathway size. Thus, pathway size may contribute to approximately 30% of the ranking metric.

To confirm that this unconnected graphlet was not negatively impacting our ranking metric, we also examined its contribution to implausible pathway detection performance. On the 3 NetPath validation pathways, we found that using only the graphlet with 4 disconnected nodes resulted in poor performance (Figure S10) and that its inclusion moderately boosted performance.

Finally, we compared the graphlet frequency distance (GFD) we selected for pathway parameter advising with two alternatives: normalized graphlet frequency distance (NGFD) and graphlet correlation distance (GCD). The choice of distance calculation affects how parameter settings are ranked. When we run multiple pathway reconstruction algorithms to reconstruct the 3 NetPath validation pathways, GFD outperforms NGFD and GCD (Figure S11). One possible explanation is that unlike GCD, GFD does not attempt to eliminate the signal of global topological properties such as size. Some signal of global topology likely helps identify which reconstructed pathways are similar to reference pathways, improving the NetPath reconstruction.

## 3 Discussion

Pathway parameter advising selects parameters that lead to useful, plausible pathways for a variety of pathway reconstruction algorithms. This parameter tuning approach is algorithm-agnostic and uses background knowledge in the form of pathway databases to succeed in selecting reasonable pathways during pathway reconstruction.

Many of the networks down-ranked by pathway parameter advising, such as pathways with many thousands of nodes or pathways consisting only of a single node and its neighbors, seem obvious to avoid. Typically, these types of reconstructed pathways are ignored through a process of manual trial and error. Any manual step in the pathway analysis could lead to human error, may accidentally introduce bias into the final pathway model, and limits the number of parameter combinations that can be assessed. Therefore, automatically avoiding these poor pathways is important. The specific choice of criteria for defining implausible pathways was inconsequential. Pathway parameter advising excelled at implausible pathway detection compared to other parameter ranking methods for all definitions of implausibility. Pathway parameter advising quantifies and deprioritizes implausible topologies without any human intervention except for the inclusion of background knowledge. In addition to avoiding implausible pathways, pathway parameter advising narrowly performed best at reconstructing NetPath pathways. Although it was less clearly dominant than in the implausible pathway detection experiment, this further highlights its effectiveness. Much of the total performance is driven by how well pathway parameter advising performs in PCSF, though it is also the only parameter selection strategy to be either the first or second highest performing for all 4 pathway reconstruction methods. However, the raw MCC values in many test cases were so low that differences in MCC were driven by only a few interactions, so this experiment alone does not provide enough evidence to draw strong conclusions.

We also found that no single graphlet dominated our ranking metric. There are likely two causes. In larger networks, such as complete protein interaction networks, disconnected graphlet counts can dominate other graphlets by orders of magnitude (Johansson *et al*., 2015). In contrast, the networks in biological pathway reconstruction tend to not be large enough for the portion of unconnected graphlets in sparse graphs to completely dominate other graphlets. In addition, our ranking metric only calculates distance from the closest 20% of reference pathways. Thus, the signal of pathway size from the 4 unconnected nodes graphlet guides the selection of the closest reference pathways to pathways of similar size. Within this 20%, other graphlets representing more local topology have a larger contribution.

Although we used different pathway databases in our experiments for reconstructing pathways from sampled nodes and the set of reference pathways, it is possible that some pathways are similar across the NetPath and Reactome databases. Cross-database pathway similarity could cause a version of the reconstructed pathway to be used as a reference pathway. However, even if this is the case, the shared pathway would be 1 of the over 1,000 Reactome pathways used as a reference. Thus, its effect on the ranking metric would be negligible.

In all experiments, other parameter selection methods especially struggled choosing parameters for PCSF. Finding a good parameter setting for a method with multiple, complex parameters like PCSF can be especially difficult and important and is where pathway parameter advising is most useful. In contrast, a method like PathLinker contains a single parameter, which monotonically increases pathway size. Changing the parameter value has a relatively predictable effect.

There are some drawbacks to pathway parameter advising. It requires a parameter sweep as opposed to a single run with the default parameter setting. This greatly increases overall runtime for some pathway algorithms. Because pathway parameter advising is algorithm-agnostic, it makes no assumptions about the parameter space it is optimizing. Thus, pathway parameter advising has no way of knowing if the range of parameters considered is broad enough to find the optimal pathway. However, it is worth noting that all other parameter selection methods tested except for the default parameters suffer from this drawback as well.

Another potential limitation is that pathway parameter advising is dependent on a database of reference pathways. The popular pathway database Reactome works well in analyses here. However, if the optimal predicted pathway is reasonable but outside the range of topologies seen among the reference pathways, it would be overlooked.

Our distance metric focuses only on topology and does not include any information about the biological context. ResponseNet (Basha *et al*., 2019) and PathLinker (Youssef *et al*., 2018) extensions consider tissue-specificity and protein localization context, respectively. A possible extension of pathway parameter advising would be to account for this information, such as adding a penalty for interactions that occur in different tissues or cellular compartments. Similarly, we used the entire Reactome database as the reference pathways. Limiting the reference pathways to a certain process or function, such as signal transduction or disease, could allow pathway parameter advising to select pathways more similar to a domain of interest. In addition, instead of computing graphlet distance using 20% of all reference pathways, we could first cluster the reference pathways and consider the distances only to pathways in the most topologically similar cluster.

Now that we have demonstrated the utility of pathway parameter advising to evaluate predicted pathways, it could be wrapped with hyperparameter optimization to fully automate the process with a Bayesian optimization framework (Kandasamy *et al*., 2019; Wang *et al*., 2013; Chen *et al*., 2012). This would allow for complete and standardized automation of the pathway reconstruction process. It could reduce the overall time required to select parameters as well because the parameter space can be explored adaptively instead of through an exhaustive grid search. Avoiding a parameter grid search is most valuable for pathway reconstruction methods like PCSF and ANAT (Yosef *et al*., 2011) that have several tunable parameters.

## 4 Methods

### 4.1 Pathway Parameter Advising

Pathway parameter advising is based on the parameter advising framework (DeBlasio and Kececioglu, 2015). A parameter advisor consists of two parts: a set of candidate parameter settings *S* and an accuracy estimator *E*. The parameter advisor evaluates each candidate parameter setting in *S* using *E* to estimate the optimal parameter set. In order to adapt parameter advising to the pathway reconstruction task, we must choose a function *E* that can estimate the quality of a reconstructed pathway. While we do not have a direct way to define what criteria an optimal solution satisfies, we do have access to pathways that match biologist intuition of what a biological pathway should look like. Curated pathway databases, such as the KEGG (Kanehisa and Goto, 2000), Reactome (Fabregat *et al*., 2018), and NetPath (Kandasamy *et al*., 2010), contain pathways that have been compiled by biologists. Therefore, we can construct our estimator around these curated pathways. This leads to the key assumption of pathway parameter advising: *reconstructed pathways more topologically similar to manually curated pathways are more useful to biologists*.

Our parameter tuning approach requires the inputs to the pathway reconstruction algorithm, a set of candidate parameter settings, and a set of pathways from a reference pathway database. Pathway reconstruction algorithms’ input typically consists of an interactome, such as STRING (Szklarczyk *et al*., 2019), and a set of biological entities of interest, such as genes or proteins. We refer to the pathways created by the algorithm as *reconstructed* pathways and the curated database pathways as *reference* pathways. Pathway parameter advising uses a graphlet distance-based estimator *E* to score each reconstructed pathway’ s similarity to the reference pathways. It uses these scores to return a ranking of the reconstructed pathways (or their respective parameter settings).

Pathway parameter advising is designed to be method-agnostic. It can be run with any pathway reconstruction algorithm that outputs pathways and has user-specified parameters. Currently, pathway parameter advising is designed to examine undirected graphs, and directed graphs are converted to be undirected.

#### 4.1.1 Graphlet Decomposition

In order to topologically compare reconstructed and reference pathways, we first decompose all pathways into their graphlet distributions. A graphlet is a subgraph of a particular size within a network. The concept of graphlets is similar to that of network motifs (Milo *et al*., 2002). However, network motifs typically refer to graphlets that appear in a network significantly more often than expected by chance.

Original work with graphlets only considered connected graphlets to better capture local topology (Pržulj *et al*., 2004). However, we use both connected and disconnected subgraphs, thus allowing all possible combinations of nodes in a pathway to be considered a graphlet. This allows our parameter ranking to capture global topological properties such as pathway size in addition to local topology. One disadvantage of disconnected graphlet counts is that the counts of disconnected graphlets, such as the graphlet containing 4 unconnected nodes, grow at a much faster rate than those of connected graphlets in sparse networks. However, this does not adversely affect our ranking metric (Section 2.5).

Pathway parameter advising uses the parallel graphlet decomposition library (Ahmed *et al*., 2015) to calculate counts of all graphlets up to size 4 in a pathway. This constitutes 17 possible graphlets (Figure 3). We convert these counts into frequencies and represent each pathway by a vector of 17 values between 0 and 1. This vector, referred to as the graphlet frequency distribution, summarizes the topological properties of a pathway, allowing us to quantify topological similarity.

#### 4.1.2 Distance Calculation

To calculate the topological distance between two pathways, we take the pairwise distance of their graphlet frequency distributions. For pathways *G* and *H*, we denote their frequencies of graphlet *i* as *F*_*i*_(*G*) and *F*_*i*_(*H*), scalars between 0 and 1. We then define the graphlet frequency distance *D*(*G, H*) as

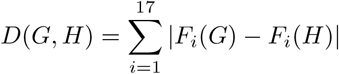

This differs from relative graphlet frequency distance, which log transforms and scales the raw graphlet counts (Pržulj *et al*., 2004). We considered other graphlet-based metrics such as a variation of relative graphlet frequency distance and graphlet correlation distance (Yaveroğlu *et al*., 2014) but found that they performed worse in our preliminary analyses (Figure S11).

#### 4.1.3 Ranking Parameters

After calculating the graphlet frequency distribution for each reconstructed and reference pathway, we can calculate their mean graphlet frequency distance to the reference pathways to get *E*. When calculating this aggregate distance, we only consider the 20% closest reference pathways to the reconstructed pathway. The threshold choice has little impact on the parameter ranking (Figure S12). It is motivated by not requiring a reconstructed pathway to be similar to every reference pathway, but instead similar to at least some reference pathways. Thus, a pathway *G*’ s score *E*(*G*) is calculated as

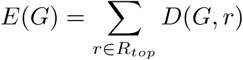

where *R*_*top*_ is the set of the 20% closest reference pathways to *G*. The pathways, or equivalently the parameters used to generate those pathways, are sorted by *E*(*G*) in ascending order. Once the final ranking is created, the top reconstructed pathway can be used for downstream analysis. Alternatively, the top *n* pathways can be merged into an ensemble pathway.

### 4.2 Pathway Reconstruction Methods

Pathway reconstruction algorithms were chosen to have a wide range of methodologies, from NetBox’ s statistical test to PathLinker’ s weighted shortest paths algorithm. We used the following 4 methods for our implausible pathway detection and NetPath reconstruction experiments. These methods and the parameters tested are summarized in Table 1.

**Table 1.**
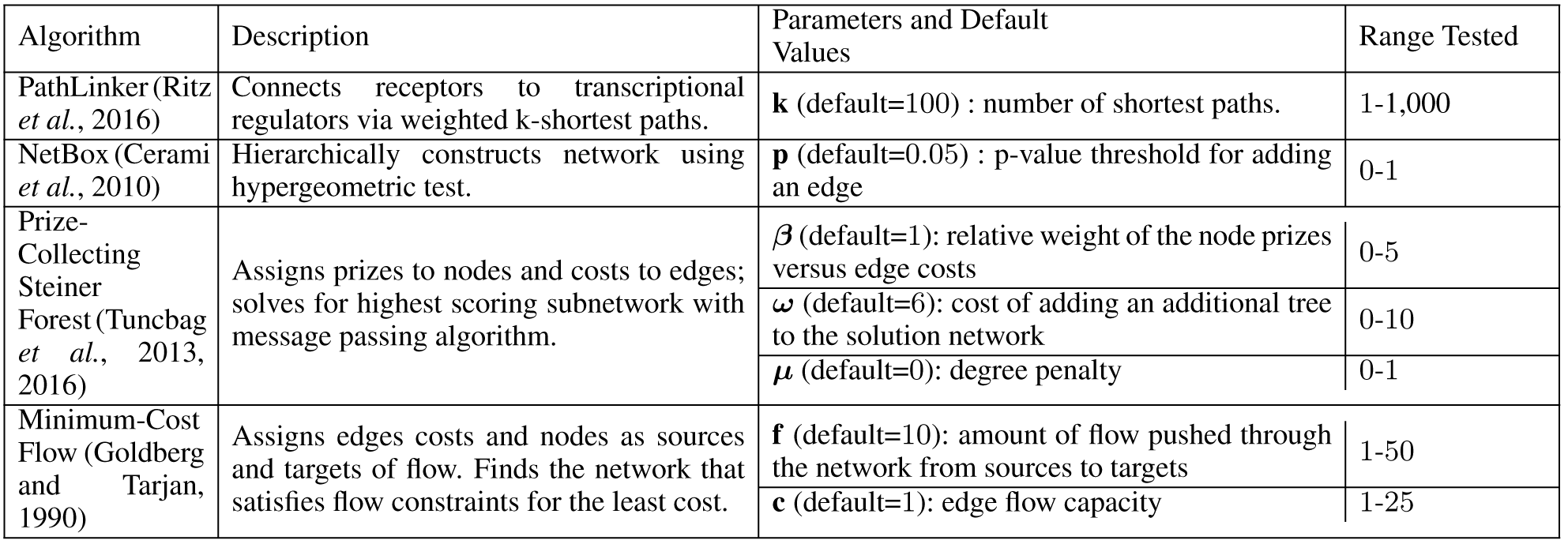
The 4 pathway reconstruction methods and the parameters tuned for each.

#### PathLinker

PathLinker (Ritz *et al*., 2016) reconstructs pathways based on a weighted *k* shortest paths algorithm. It finds paths between sets of receptors and transcriptional regulators, similar to the source and target nodes in minimum-cost flow. It is controlled by the parameter *k*, which defines how many paths to return in the final network. We varied *k* from 1 to 1,000 in increments of 1. We used PathLinker version 1.1 for all analyses.

#### NetBox

NetBox (Cerami *et al*., 2010) hierarchically constructs networks from a set of input nodes. At each iteration, it searches for linker nodes that connect two nodes in the current network. It then chooses to add these linker nodes to the network based on the results of a hypergeometric statistical test comparing the degree of the linker node to how many nodes in the pathway it connects. NetBox is controlled by the parameter *p*, a p-value threshold, which sets the threshold for whether or not linker nodes should be included. We varied *p* from 0 to 1 on a log scale from 1*×* 10^−30^ to 1 in increments of half an order of magnitude, giving a total of 60 steps. We used NetBox version 1.0 for all analyses.

#### Prize-Collecting Steiner Forest

In PCSF (Tuncbag *et al*., 2013, 2016; Kedaigle and Fraenkel, 2018), nodes are assigned prizes and edges are given costs. The optimal subnetwork, which is found via a message-passing algorithm (Bailly-Bechet *et al*., 2011), is the pathway *F* consisting of nodes *N*_*F*_ and edges *E*_*F*_ that best balances collected prizes versus cumulative edge costs according to the following function:

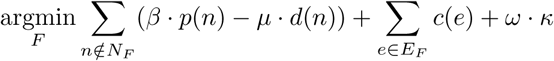

where *p*() is the positive prize for each node, *d*() is a node’ s degree, *c*() is the cost of each edge, and *κ* is the number of connected components in the pathway. The optimal subnetwork is always a tree- or forest-structured graph. We varied 3 PCSF parameters: *β*, which controls the relative weight of the node prizes versus edge costs was varied from 0 to 5 in increments of 0.5; *µ*, which affects the penalty for high-degree nodes was varied from 0 to 1 in increments of 0.1; and *ω*, which controls the cost of adding an additional tree to the solution network was varied from 0 to 10 in increments of 1. We used version 1.3 of the msgsteiner message-passing algorithm and version 0.3.1 of OmicsIntegrator for all analyses.

#### Minimum-Cost Flow

The minimum-cost flow problem assigns certain nodes in the network to be “sources” and others to be “targets”. Edges, which transport the flow from node to node, have a cost associated with using them and a capacity of how much flow they can hold. The solution is the network that satisfies the flow requirements of the source and target nodes while using lowest cost in edges (Goldberg and Tarjan, 1990). We implemented a version of min-cost flow using the solver provided in Google’ s OR-Tools^1^, which solves the min-cost flow problem using the algorithm outlined in Bünnagel *et al*. (1998). This is a generic version of the algorithm used in ResponseNet (Basha *et al*., 2013). Two parameters control the min-cost flow solution: the total flow through the network, which we vary from 1 to 50 in increments of 1, and the edge flow capacity, which we vary from 1 to 25 in increments of 1. We used Google’ s OR-Tools version 7.1.6720 for all analyses.

### 4.3 Alternate Parameter Selection Methods

We compare our pathway parameter advising approach to the following parameter selection strategies from the literature:

#### Cross-validation

CV involves splitting the input data, the biological nodes of interest provided to the pathway reconstruction algorithm, into training and testing sets multiple times for each parameter setting. For example, the input data could be sampled nodes from a NetPath pathway. A pathway reconstruction method is then run on each training set and evaluated on each respective testing set. In this problem setting, we do not have external ground truth with which to evaluate the predictions on test set data. Instead, we perform 5-fold CV on subsets of the input data, producing a pathway from the training set nodes. The parameter values that produce pathways that recover the highest proportion of the test set nodes are chosen.

#### ResponseNet ranking

We also tested a parameter selection heuristic used by ResponseNet (Yeger-Lotem *et al*., 2009). The criterion is to select parameters that result in a pathway whose nodes include at least 30% of the input data, while having the lowest proportion of low confidence edges. We extend this to rank the pathways that do include 30% of the inputs by their proportion of low confidence edges, followed by the pathways that include less than 30% of the inputs to form a full ranking.

#### PCSF Robustness Ranking

As suggested by Kedaigle and Fraenkel (2018), for PCSF we can also rank pathways by their robustness. Robustness is measured by how often nodes appeared in multiple runs with small random perturbations to the scores on the input nodes. We only applied this strategy to reconstructed pathways from PCSF. Although it could be adapted to other pathway reconstruction methods, we decided to use it only in the method for which it was directly implemented.

### 4.4 Evaluating Reconstructed Pathway Plausibility

In order to examine the ability of pathway parameter advising to avoid parameter settings that lead to impractical pathways, we created topological criteria that we use to define pathways as plausible or implausible. These criteria are based on the literature where possible and were created without considering the topology of pathways from pathway databases. However, given that curated pathway databases are also based on information from the literature, these plausibility criteria should not be considered completely independent from the pathway databases. We use these criteria as a heuristic to label pathways as positive (plausible) or negative (implausible). The labels enable us to evaluate pathway rankings as a classification problem, determining if a method can correctly rank plausible reconstructed pathways before implausible pathways. These criteria are based on previous analyses of biological networks, and are as follows:

#### Size

We allowed pathways that had between 10 and 1,000 nodes. Pathways whose size was outside this range are not practical for hypothesis generation and downstream analysis.

#### Hub node dependence

A common issue with pathway reconstruction is an over-reliance on high-degree or hub nodes. Dominant hub nodes can create pathways consisting almost entirely of a single node and its neighbors with few to no connections between those neighbors (Kedaigle and Fraenkel, 2018). We score hub node dependence using the ratio of the degree of the highest degree node to the average node degree of the entire pathway. If the maximum degree is more than 20 times greater than the average degree, we consider the pathway implausible.

#### Clustering coefficient

Biological networks have been found to have clustering or community structure that is hierarchical (Ravasz *et al*., 2002); communities within the network exist at multiple scales and are often nested within each other. Thus, it would be reasonable to expect a plausible biological pathway to have at least a moderate level of community structure. We calculate the average clustering coefficient of all nodes in the pathway, a common metric for measuring community structure (Barabási and Oltvai, 2004). The clustering coefficient of a node is the proportion of its neighbors that are also neighbors of each other. This can be averaged over all nodes in the pathway as a measure of the overall level of clustering. We require pathways to have an average clustering coefficient of at least 0.05, as we expect at least some small level of clustering to exist. Because this requirement eliminated all PCSF pathways in 25% of tasks, when evaluating PCSF we excluded this metric in all cases.

#### Assortativity

A network’ s level of assortative mixing is defined as the tendency of high-degree nodes to be connected to other high-degree nodes. Biological networks have been found to be generally dissasortative, meaning that high-degree nodes tend to be connected to low-degree nodes (Newman, 2002; Albert *et al*., 2011). Assortativity is measured between−1 and 1, where assortative networks have positive values and dissasortative networks have negative values. This value can be viewed as the correlation between a node’ s degree and its neighbors’ degrees within the pathway. We consider pathways with assortativity between −1 and 0.1 plausible to allow for some leeway in pathways being slightly assortative.

We selected these criteria based on attributes it would be reasonable to expect a biological pathway to have, with values supported by the literature where possible. If any of these criteria are not met, we consider the pathway to be implausible. These thresholds were not influenced by the graph topologies in the reference pathway database in order to minimize circularity between the reference pathway-based rankings and the plausibility criteria used to evaluate those rankings.

However, most of the reference pathways we considered happen to be plausible. 77% of the Reactome pathways we used as reference pathways are plausible, though it should be noted that these Reactome pathways have already been filtered by size (Section 4.6.2).

While the criteria for defining a plausible network are useful for comparing networks created by the same method with different parameter settings, they should not be considered as a metric for comparing pathways across pathway reconstruction methods. Different pathway reconstruction methods are able to use different sources of information and have complex strengths and weaknesses beyond the local topologies they return. For instance, NetBox, which had the highest proportion of plausible pathways, cannot take into account information such as edge confidence or scores on proteins of interest that other methods such as PCSF can. Furthermore, the 4 plausibility properties are a binary way to determine if a pathway is reasonable or unreasonable. They cannot be used to rank pathway reconstruction parameters.

In order to make sure that our experimental results are not overly sensitive to the specific choice of thresholds for pathway plausibility, we tested other thresholds in a grid search. We varied the maximum network size threshold from 200 to 2,000, the hub node dependence measure from 5 to 50, the clustering coefficient threshold from 0.0 to 0.1, and the assortativity threshold from −0.5 to 0.5. Each range was divided into 10 intervals, for a total of 10,000 sets of plausibililty thresholds. Figures 5 and S1 evaluate the pathway reconstruction methods across these different thresholds.

### 4.5 Evaluating NetPath Pathway Reconstructions

When comparing reconstructed pathways to the original NetPath pathways in Section 2.3, we used MCC to quantify the reconstruction quality (Matthews, 1975). MCC is a metric used in binary classification that ranges between −1 and 1, where 1 indicates a perfect binary classification and −1 indicates a completely incorrect classification. It can be viewed as the correlation between the predicted and true labels in a classification task. MCC has been shown to be well suited to evaluate classification in imbalanced settings (Boughorbel *et al*., 2017). In order to treat comparing a reconstructed and a NetPath pathway as a classification task, we consider all edges in a NetPath pathway as the positive set and all other edges as the negative net. MCC is then defined as

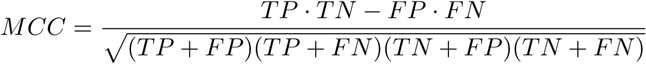

where TP is the number of true positives (edges that appear in both the NetPath and reconstructed pathway), FP is the number of false positives (edges that do not appear in the NetPath pathway but do in the reconstructed pathway), TN is the number of true negatives (edges that do not appear in either the NetPath pathway or the reconstructed pathway), and FN is the number of false negatives (edges that appear in the NetPath pathway but not in the reconstructed pathway). When comparing MCC values of multiple pathways and reconstruction methods, we normalized MCC values by the best possible MCC among all tested parameter values for that pathway and method. We refer to this value as the adjusted MCC.

### 4.6 Datasets

#### 4.6.1 Interactomes

For both PathLinker and NetBox, we used the interactome included as a part of their software packages. For PCSF and min-cost flow, we used an interactome from Köksal *et al*. (2018) that merged protein interactions from the iRefIndex database v13 (Razick *et al*., 2008) and kinase-substrate interactions from PhosphoSitePlus (Hornbeck *et al*., 2014). The interactions from the iRefIndex database include confidence scores, while confidence scores for kinase-substrate interactions were inferred from the number of interactions for each kinase-substrate pair and the type of experiment that detected the interaction. If an interaction was included in both databases, the PhosphoSitePlus interaction was used. This resulted in a network with 161,901 weighted edges.

#### 4.6.2 Pathway Databases

All parameter tuning was performed with Reactome as the set of reference pathways. Reactome (Fabregat *et al*., 2018) is a database of manually curated pathways, including 2,287 human pathways. Reactome is open-source, where all contributions must provide literature evidence and are reviewed by an external domain expert before being added. Pathways smaller than 15 nodes were excluded as too small for meaningful interpretation. Reactome pathways were retrieved using Pathway Commons (Rodchenkov *et al*., 2019). Pathway Commons converted from the Reactome data model to a binary interaction model using a set of rules for each interaction type ^2^.

The implausible pathway detection and NetPath reconstruction experiments were performed on pathways from the NetPath database. NetPath is a collection of 36 manually curated human signal transduction pathways (Kandasamy *et al*., 2010). We used 15 NetPath pathways that contain at least 1 receptor and transcriptional regulator and are sufficiently connected, as described by Ritz *et al*. (2016). We designated 3 of these NetPath pathways as validation pathways: Wnt, TGF beta, and TNF alpha. Validation pathways were used to guide the choice of distance measure. The remaining 12 pathways were reserved as test pathways for quantitative evaluations. We sampled the NetPath pathways in different ways for each pathway reconstruction algorithm to provide inputs in their expected formats, generally following the node sampling protocol that PathLinker (Ritz *et al*., 2016) used to reconstruct NetPath pathways. PCSF and NetBox do not require sources and targets, so we randomly sampled 30% of the pathway nodes as input. We also assigned each input a random prize sampled uniformly between 0 and 5 for PCSF. For PathLinker and min-cost flow, which require sources and targets, we selected all transcription factors and receptors for each pathway as outlined by Ritz *et al*. (2016).

#### 4.6.3 Influenza Host Factors

Influenza host factor data was gathered from a meta-analysis of 8 RNAi studies (Tripathi *et al*., 2015). The meta-analysis used the raw RNAi screen data to calculate a consolidated Z score for a total of 1,257 host factor genes.

### 4.7 Enrichment Analyses

GO (The Gene Ontology Consortium, 2018) and KEGG pathway (Kanehisa and Goto, 2000) enrichment was carried out with DAVID v6.7 (Huang *et al*., 2008). Enrichment was performed using GO biological process terms and all KEGG pathways. Thresholds for term inclusion were set to a count of 2 and an EASE score of 0.1.

### 4.8 Pathway Parameter Advising Implementation

A Python implementation of pathway parameter advising is available at https://github.com/gitter-lab/pathway-parameter-advising under the MIT license. While v0.1.0 and later versions of the pathway parameter advising software support Python v3.6, the results here used Python v2.7.16 and Anaconda v2019.03. The following package versions were used: pandas v0.24.2, networkx v2.2, numpy v1.16.2, matplotlib v2.2.3, and seaborn v0.9.0. The Parallel Graphlet Decomposition library was pulled from GitHub on April 30, 2019.

## 5 Data Availability

All pathway data is publicly available at http://netpath.org/browse and from https://www.pathwaycommons.org/ using the query https://www.pathwaycommons.org/pc2/search?q=*&type=pathway&datasource=reactome. The pathway parameter advising software is available from GitHub at https://github.com/gitter-lab/pathway-parameter-advising and PyPI at https://pypi.org/project/pathwayParameterAdvising/. The software is archived on Zenodo at https://doi.org/10.5281/zenodo.3985899.

## Acknowledgements

We thank Anna Ritz for providing formatted data and assisting with the NetPath evaluation, Paul Ahlquist and Eunju Park for discussing topological features of plausible pathways, Tijana Milenkovic for graphlet distance suggestions, Emma Graham Linck for testing and improving the pathway parameter advising software, and members of the Gitter research group for their helpful comments.

## Author Contributions

CM and AG developed the methods and experiments, analyzed results, and wrote the manuscript. CM wrote the software and performed the experiments. Both authors read and approved the final manuscript.

## Supplementary Information

### S1 Supplementary Figures

**Fig. S1.**
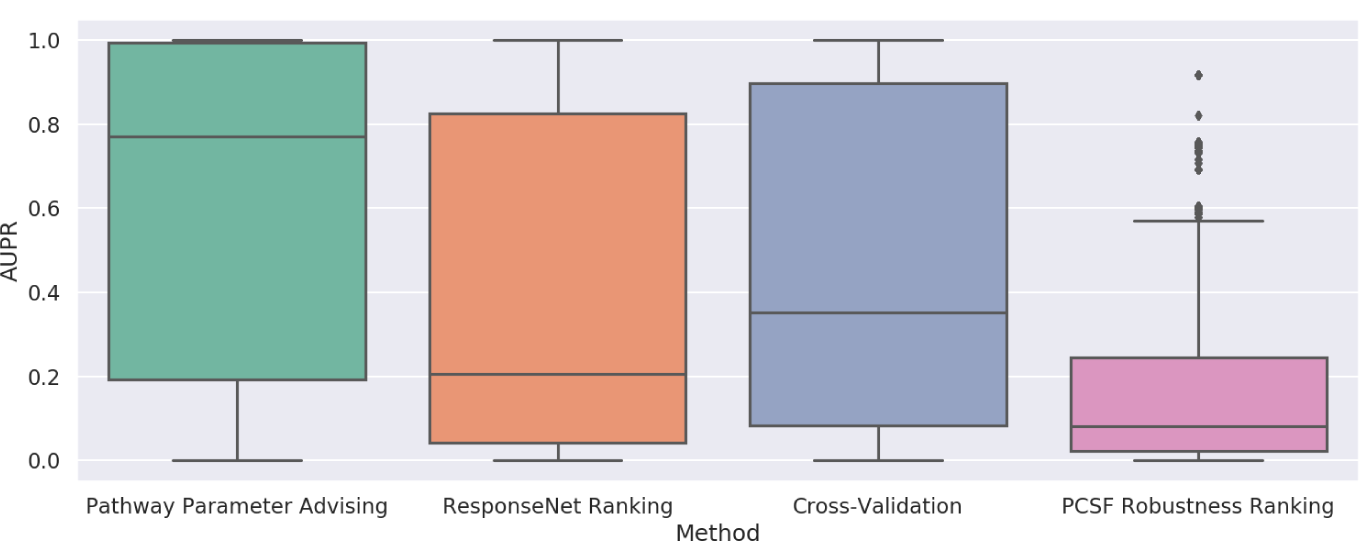
Performance of parameter selection methods on avoiding implausible networks aggregated for all considered 10,000 plausibility criteria. AUPR is shown for all 12 NetPath test pathways and 4 pathway reconstruction methods.

**Fig. S2.**
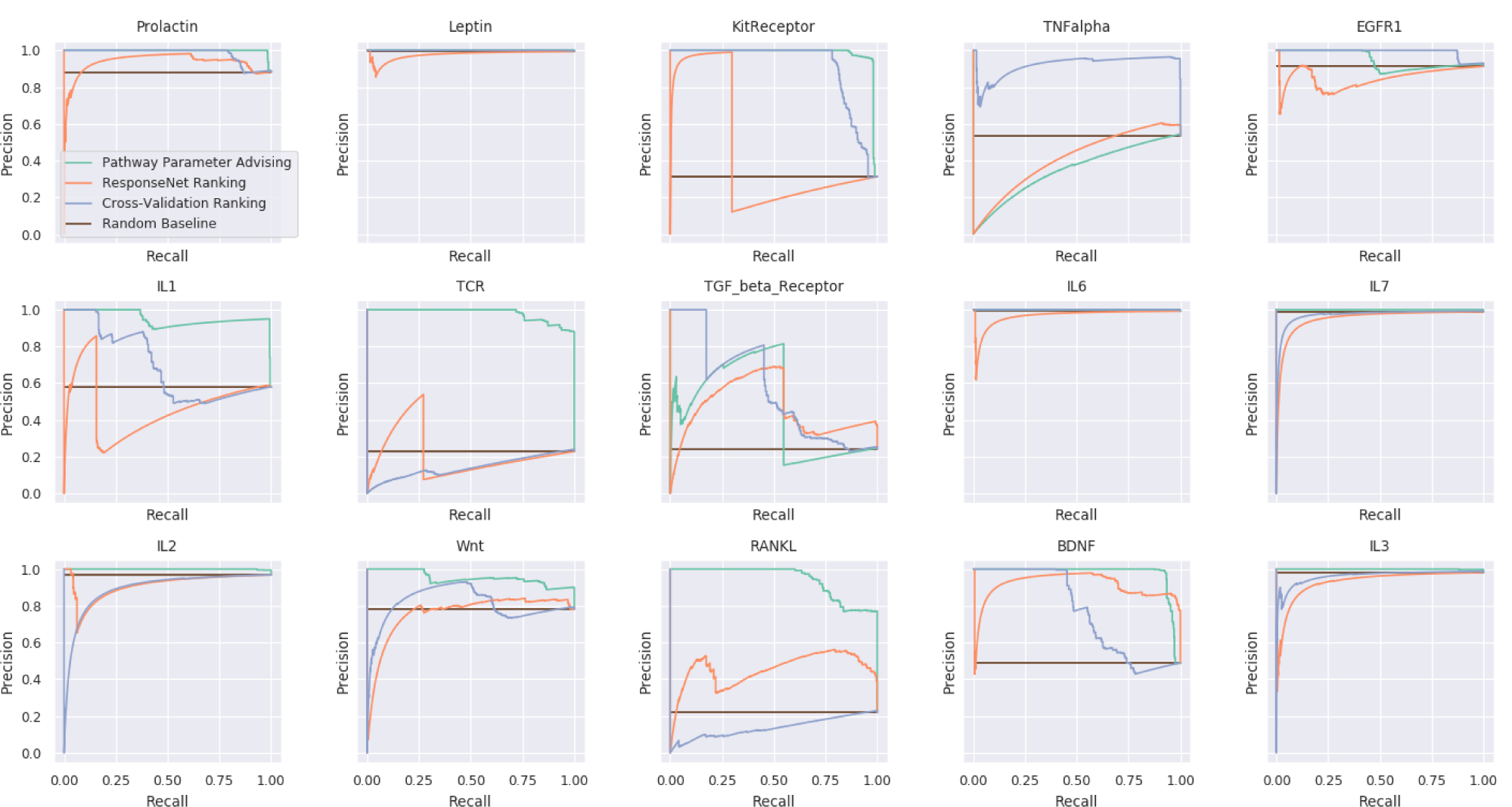
All PR curves for implausible pathway detection using different parameter ranking schemes for pathways created using PathLinker.

**Fig. S3.**
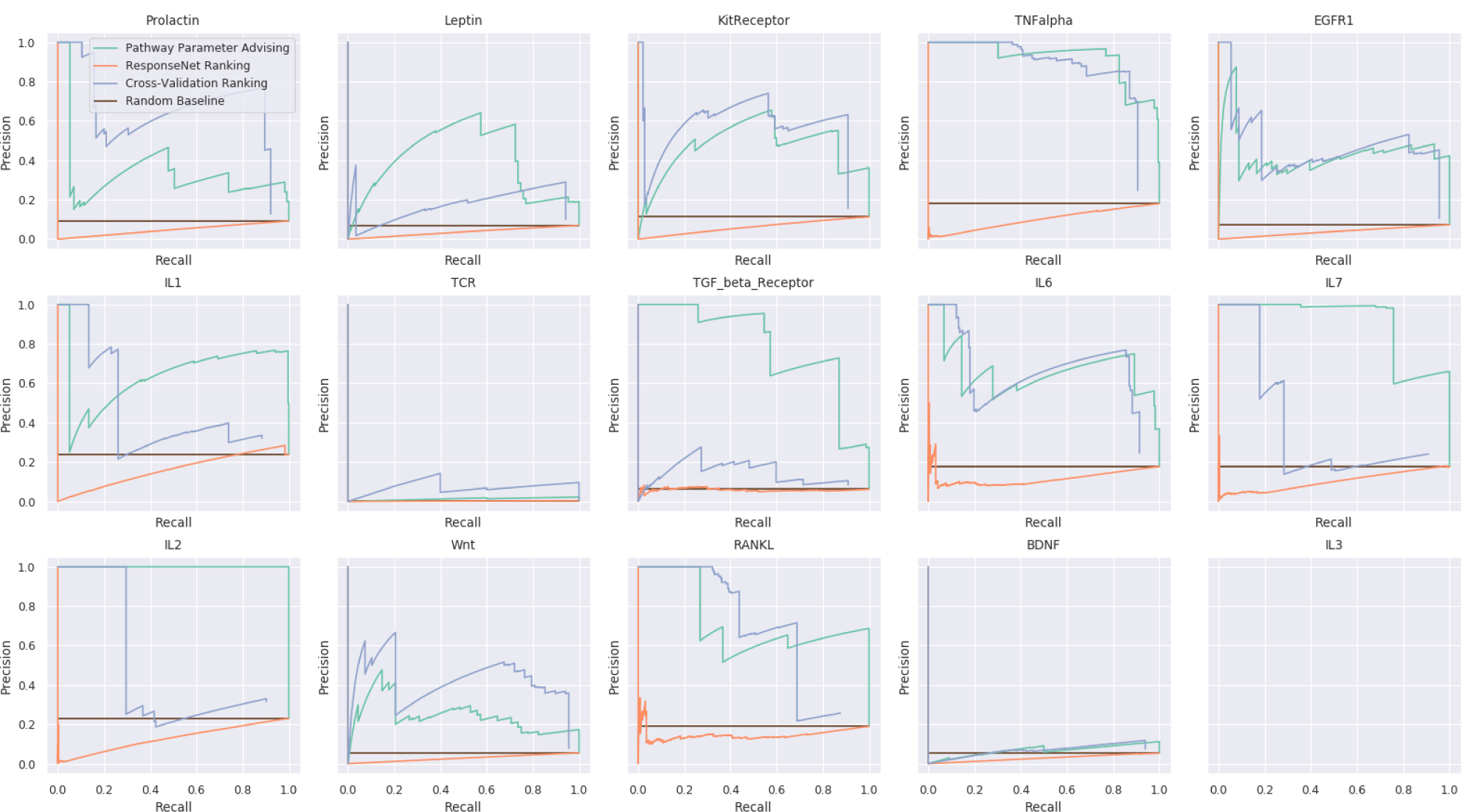
All PR curves for implausible pathway detection using different parameter ranking schemes for pathways created using minimum-cost flow. The blank panel for IL3 indicates that no reconstructed pathways were plausible.

**Fig. S4.**
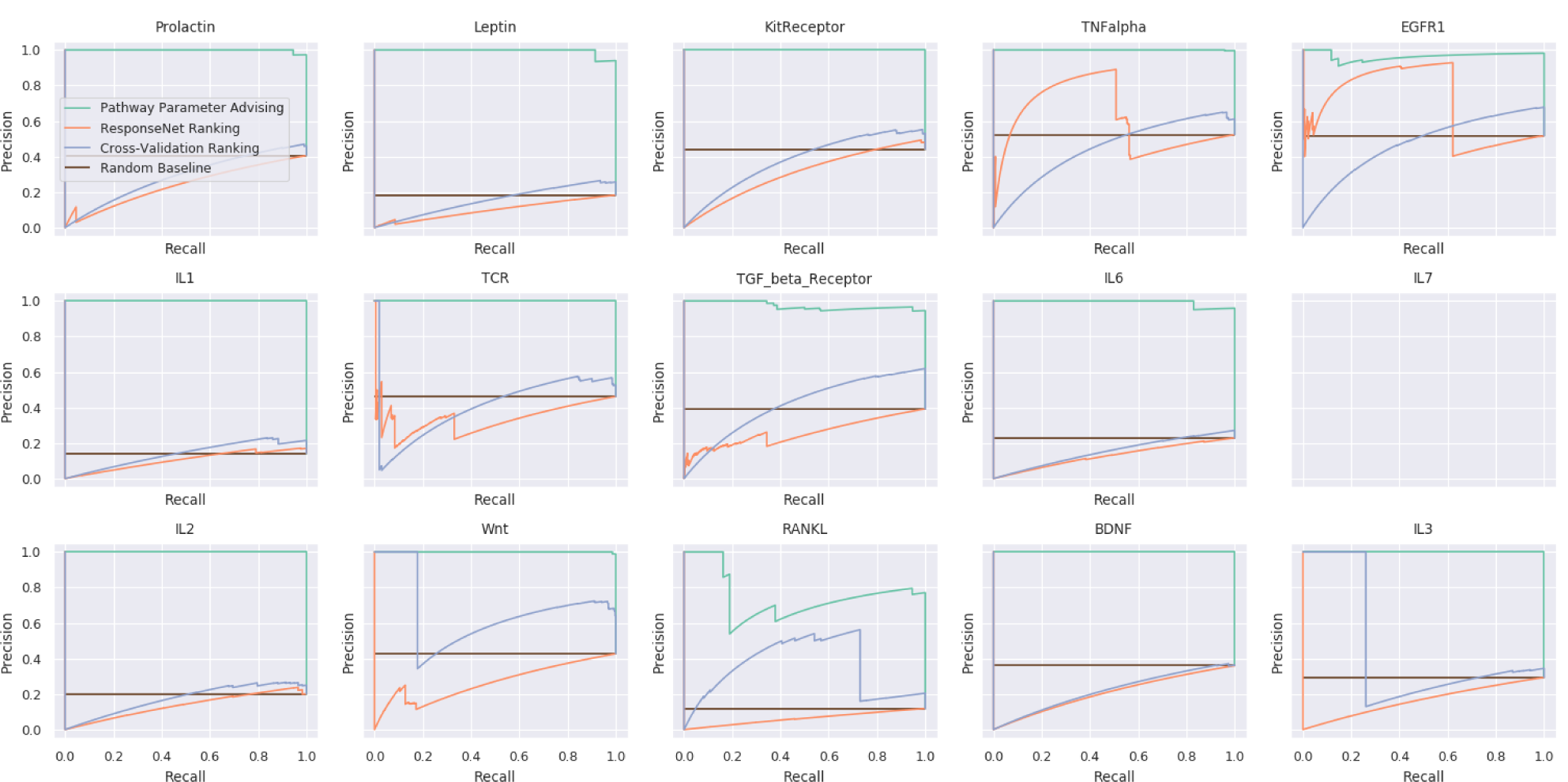
All PR curves for implausible pathway detection using different parameter ranking schemes for pathways created using PCSF. The blank panel for IL7 indicates that no reconstructed pathways were plausible.

**Fig. S5.**
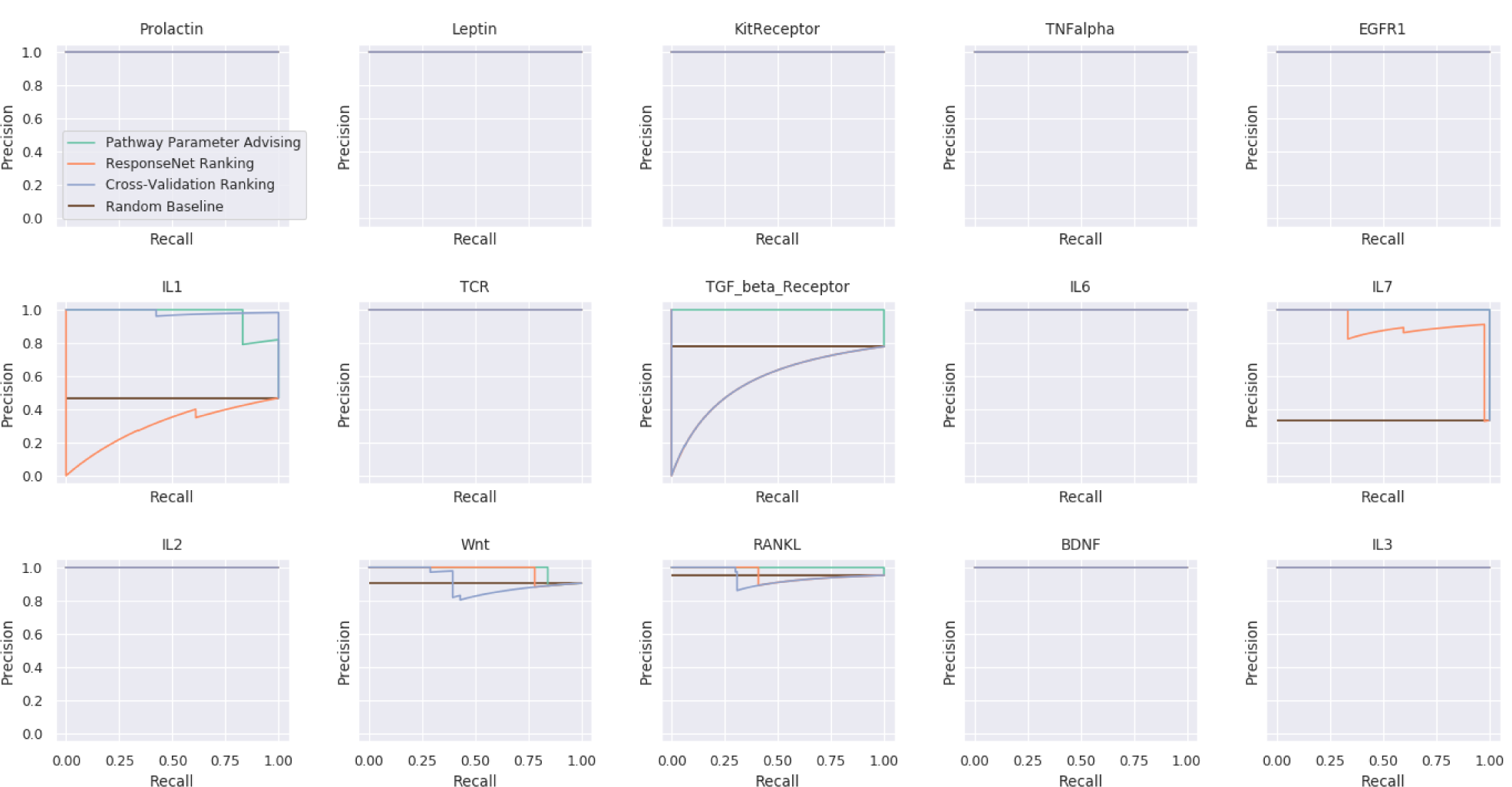
All PR curves for implausible pathway detection using different parameter ranking schemes for pathways created using NetBox. In 10 cases every reconstructed pathway met the plausible pathway criteria.

**Fig. S6.**
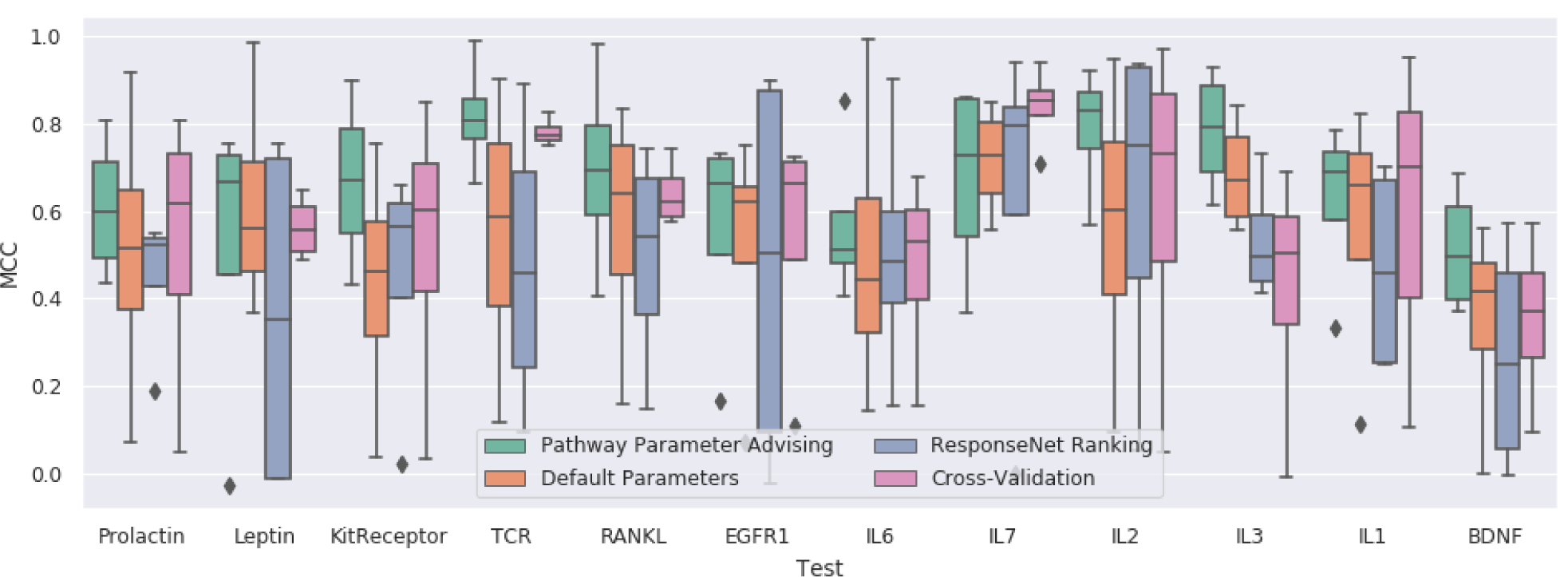
Adjusted MCC values for different parameter selection methods in the NetPath pathway reconstruction.

**Fig. S7.**
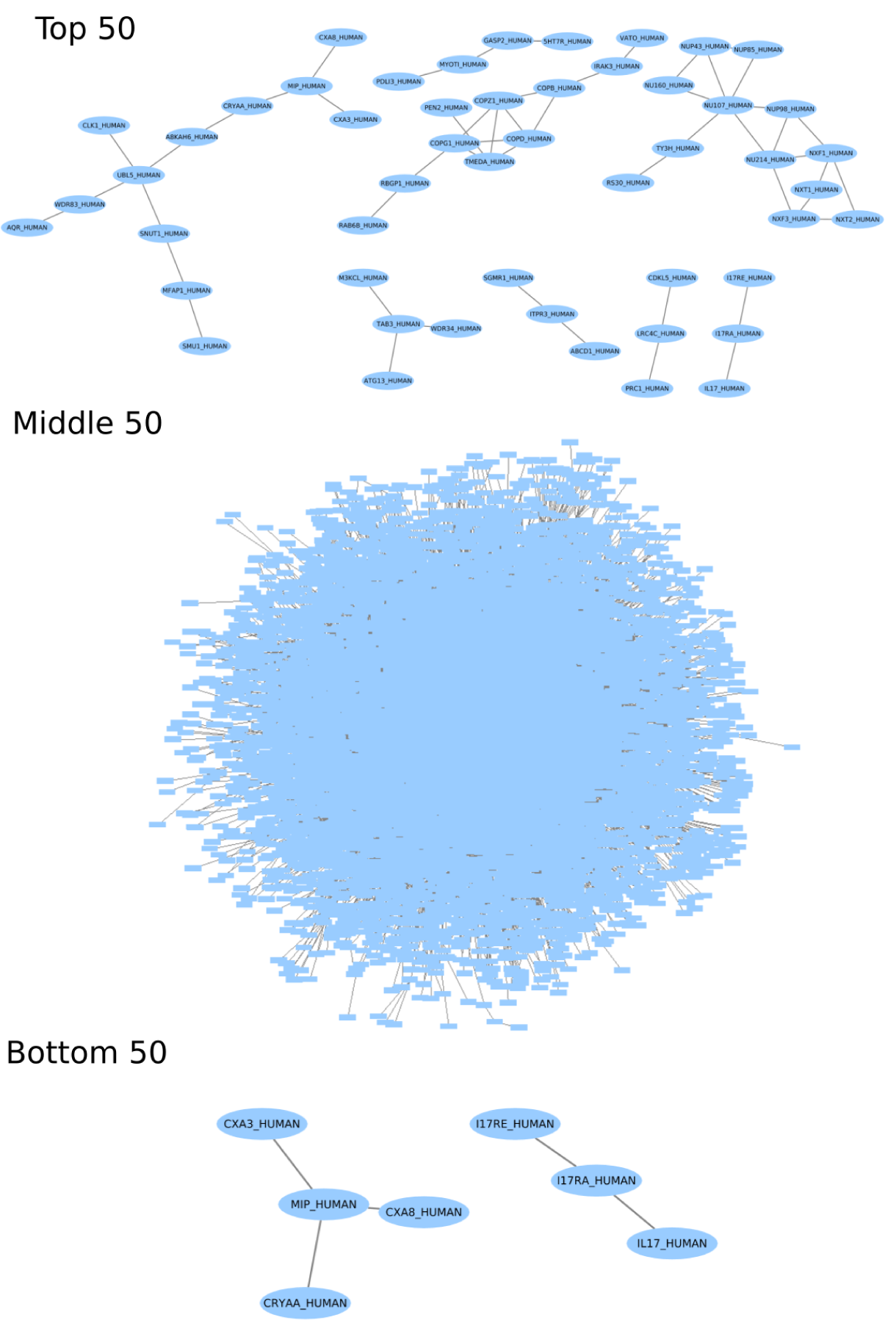
Influenza host factor pathways created by ensembling PCSF runs. The resulting pathways from the top 50, middle 50, and bottom 50 parameter settings as ranked by pathway parameter advising. All connected components over 3 nodes are shown.

**Fig. S8.**
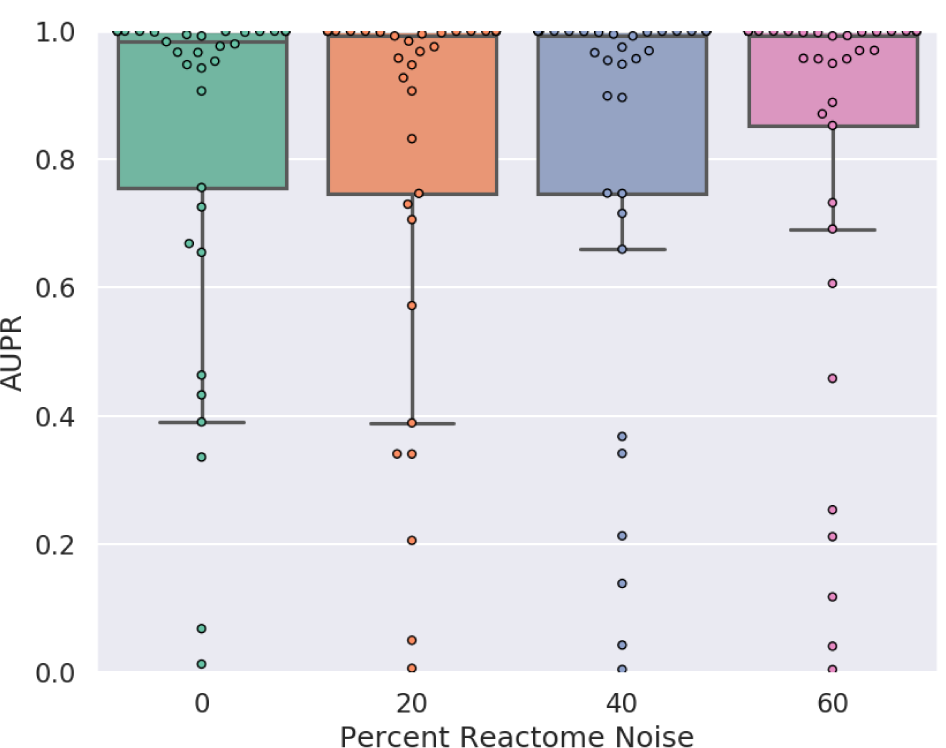
Performance of pathway parameter advising when noise is added to reference pathways. Noise was added to all Reactome pathways as described in Section 2.5, and AUPR was calculated for implausible pathway detection as described in Section 2.2 for 12 NetPath test pathways. Adding noise to the set of reference pathways resulted in almost no change to the distribution of AUPR scores over all tests.

**Fig. S9.**
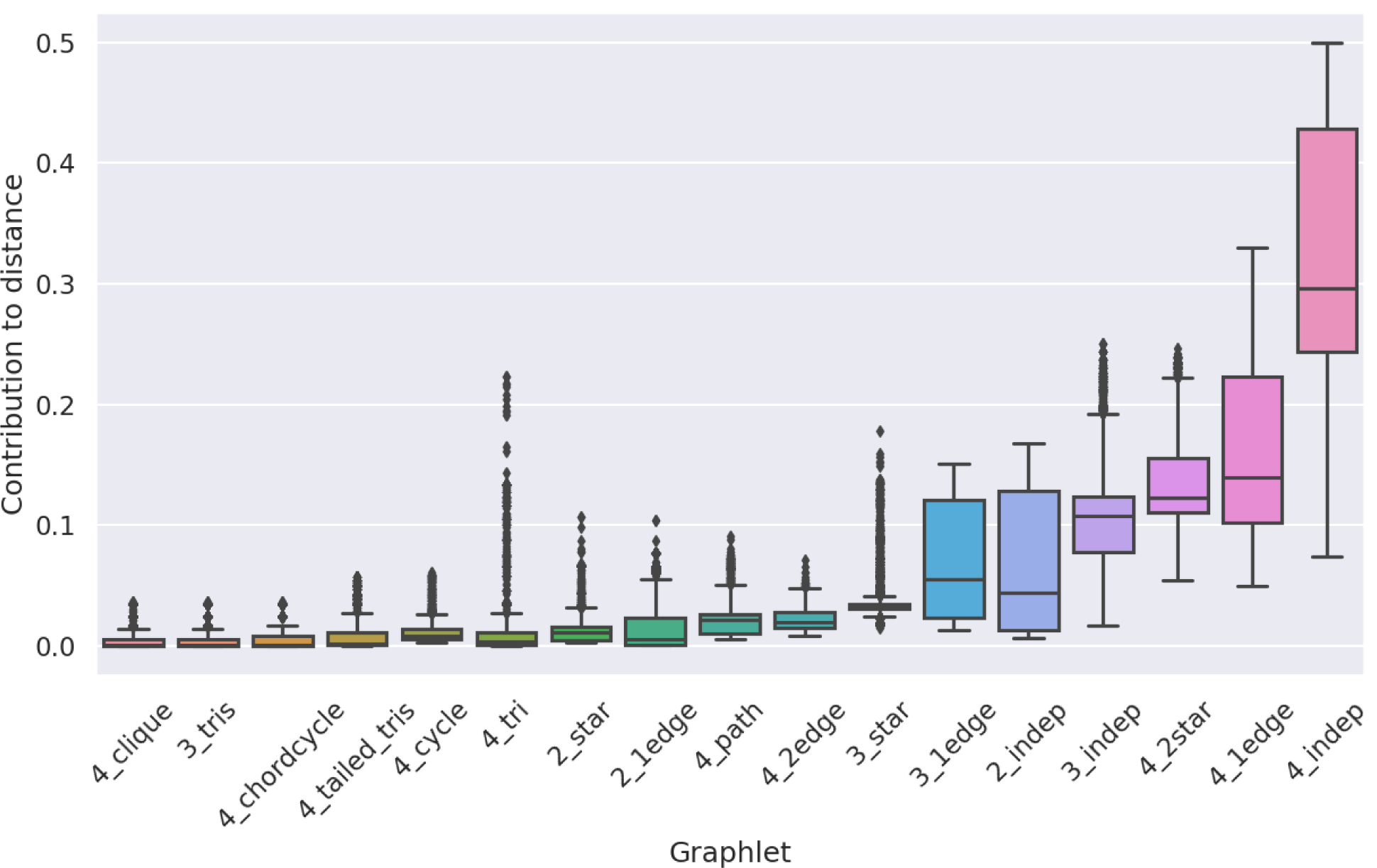
Contribution of each of the 17 graphlets to graphlet distance across the NetPath validation pathways Wnt, TNF alpha, and TGF beta and 4 pathway reconstruction algorithms. Graphlets are labeled as according to Ahmed et al. (2015). The 4 disconnected nodes graphlet, 4_indep, has a median contribution of about 30% of the GFD.

**Fig. S10.**
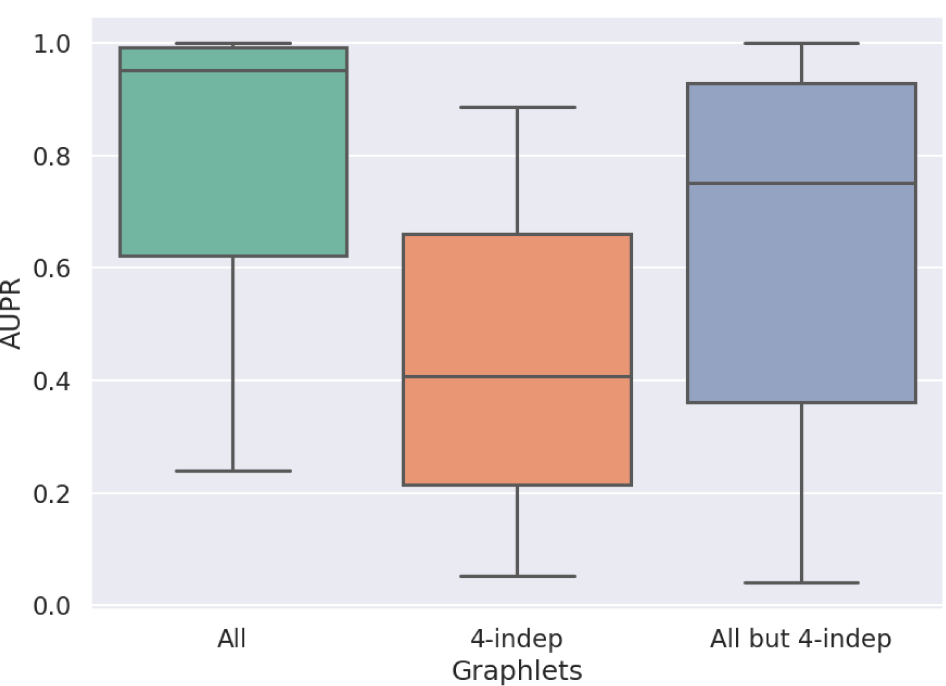
Performance of parameter selection methods on avoiding implausible networks using different sets of graphlets: all graphlets (All), only the 4 disconnected node graphlet (4-indep), and all graphlets except the 4 independent node graphlet (All but 4-indep). Boxplots represent the distributions of the AUPRs aggregated for 4 pathway reconstruction methods and the NetPath validation pathways Wnt, TNF alpha, and TGF beta. Using all graphlets yields the best performance. Removing the 4 disconnected node graphlet slightly lowers the AUPR, and only using the 4 disconnected node graphlet results in a large performance decrease. This suggests that our ranking method is highly dependent on graphlets other than the 4 disconnected node graphlet.

**Fig. S11.**
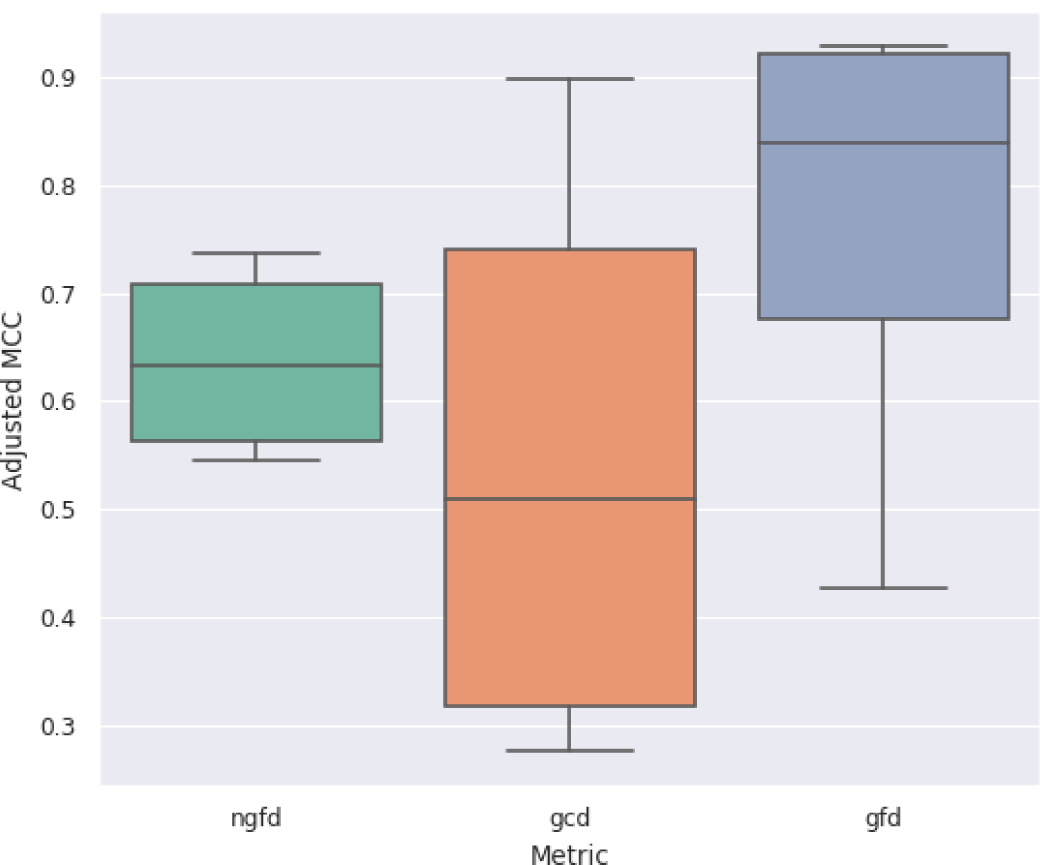
Examining the effect of different graphlet-based distance metrics on the adjusted MCC of pathway reconstruction. Reconstructions were performed on the NetPath validation pathways Wnt, TNF alpha, and TGF beta across the 4 pathway reconstruction algorithms. We examined 3 graphlet-based metrics for pathway parameter advising: normalized graphlet frequency distance (NGFD), graphlet correlation distance (GCD), and graphlet frequency distance (GFD). For NGFD, we wanted to explore a metric that takes advantage of all reconstructed pathways being sub-networks of the same interactome. Thus, we normalized all graphlet frequencies by the corresponding graphlet’ s frequency in the interactome. We also explored GCD, which measures the correlation between connected graphlets in a pathway (Yaveroğlu et al., 2014). This creates a metric that is solely focused on local topology and has minimal information about pathway size or other global topological properties. Adjusted MCC was calculated the same way as in Section 2.3.

**Fig. S12.**
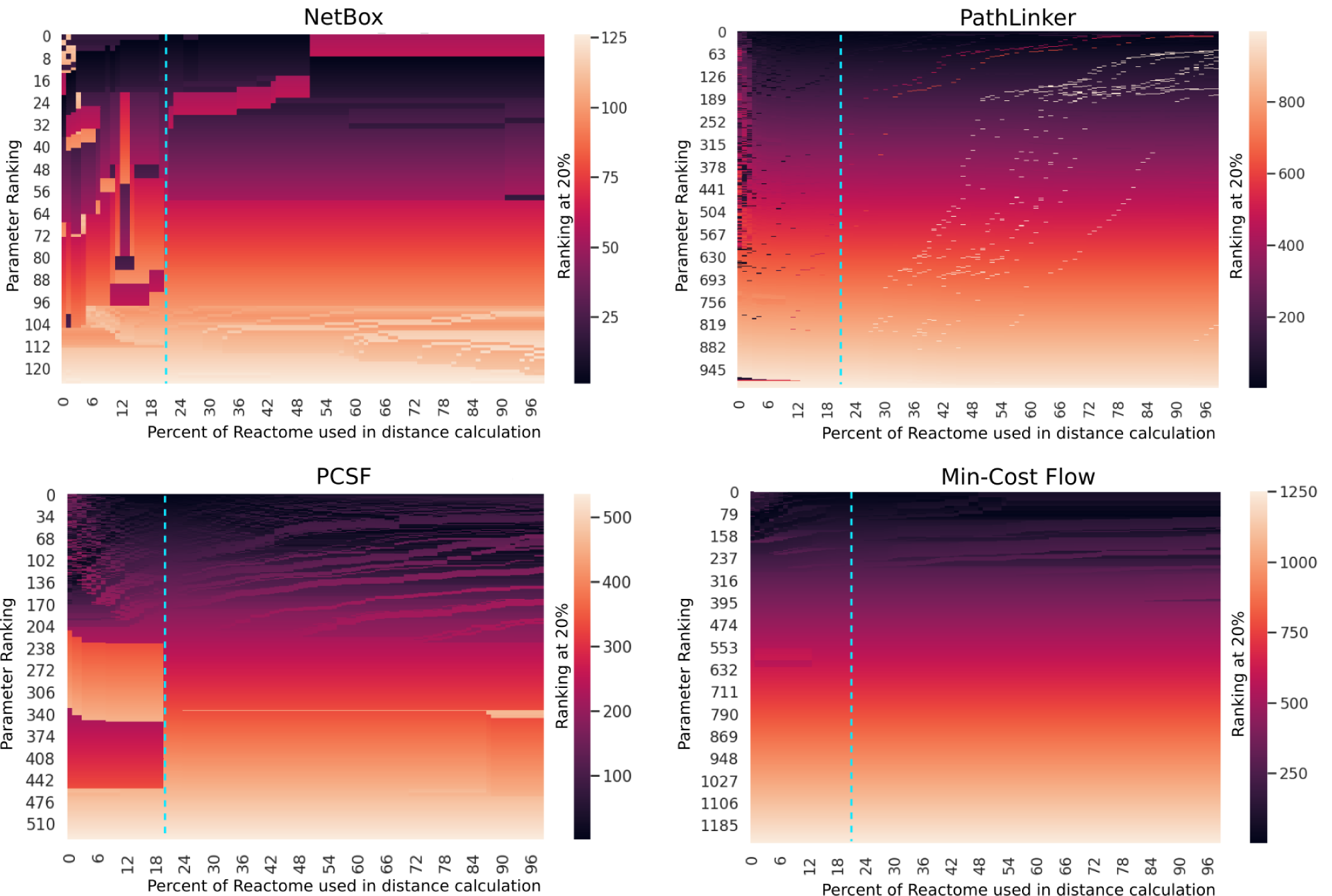
Change in ranking order over different percent thresholds for calculating score in pathway parameter advising on the TGF beta receptor NetPath pathway. Heatmaps are colored based on the ranking at the chosen threshold, 20% (marked by a blue dashed line). While the parameter ranking is unstable for very small values, by 20% the ranking remains generally unchanged as the threshold continues to increase, as can be seen in the color gradient consistency in the right halves of the figures.

### S2 Supplementary Tables

**Table S1.**
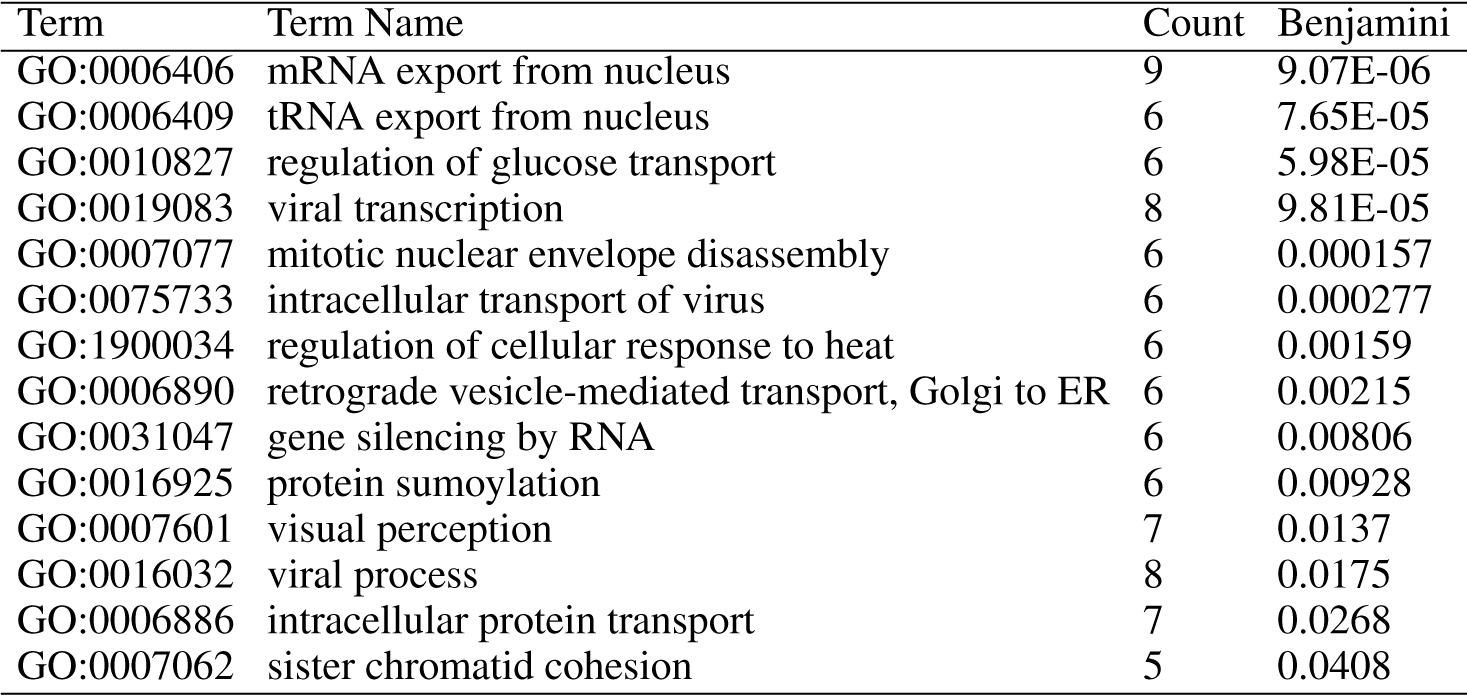
All GO terms returned from GO-term enrichment with adjusted p-value *<* 0.05 using DAVID on all nodes in the influenza host factor pathway. The influenza host factor pathway was created using the top 50 pathways ranked by pathway parameter advising constructed using PCSF. The column Benjamini shows p-values corrected using the Benjamini-Hochberg procedure.

**Table S2.** The 9 KEGG pathways returned from enrichment using DAVID of all nodes in the influenza host factor pathway. The influenza host factor pathway was created using the top 50 pathways ranked by pathway parameter advising constructed using PCSF. The column Benjamini shows p-values corrected using the Benjamini-Hochberg procedure.

| Term | Term Name | Count | Benjamini |
| --- | --- | --- | --- |
| hsa03013 | RNA transport | 10 | 4.00E-05 |
| hsa05164 | Influenza A | 6 | 0.134 |
| hsa03008 | Ribosome biogenesis in eukaryotes | 4 | 0.346 |
| hsa05323 | Rheumatoid arthritis | 4 | 0.280 |
| hsa03015 | mRNA surveillance pathway | 4 | 0.250 |
| hsa05110 | Vibrio cholerae infection | 3 | 0.433 |
| hsa00190 | Oxidative phosphorylation | 4 | 0.427 |
| hsa04721 | Synaptic vesicle cycle | 3 | 0.455 |
| hsa05120 | Epithelial cell signaling in Helicobacter pylori infection | 3 | 0.454 |

### Footnotes

1 https://developers.google.com/optimization/flow/mincostflow

2 https://www.pathwaycommons.org/pc/sif_interaction_rules.do

